# Acute multi-level response to defective *de novo* chromatin assembly in S-phase

**DOI:** 10.1101/2024.03.22.586291

**Authors:** Jan Dreyer, Giulia Ricci, Jeroen van den Berg, Vivek Bhardwaj, Janina Funk, Claire Armstrong, Vincent van Batenburg, Chance Sine, Michael A. VanInsberghe, Richard Marsman, Imke K. Mandemaker, Simone di Sanzo, Juliette Costantini, Stefano G. Manzo, Alva Biran, Claire Burny, Moritz Völker-Albert, Anja Groth, Sabrina L. Spencer, Alexander van Oudenaarden, Francesca Mattiroli

## Abstract

Long-term perturbation of *de novo* chromatin assembly during DNA replication has profound effects on epigenome maintenance and cell fate. The early mechanistic origin of these defects is unknown. Here, we combine acute degradation of Chromatin Assembly Factor 1 (CAF-1), a key player in *de novo* chromatin assembly, with single-cell genomics, quantitative proteomics, and live-microscopy to uncover these initiating mechanisms in human cells. CAF-1 loss immediately slows down DNA replication speed and renders nascent DNA hyper-accessible. A rapid cellular response, distinct from canonical DNA damage signaling, is triggered and lowers histone mRNAs. As a result, histone variants usage and their modifications are altered, limiting transcriptional fidelity and delaying chromatin maturation within a single S-phase. This multi-level response induces a cell-cycle arrest after mitosis. Our work reveals the immediate consequences of defective *de novo* chromatin assembly during DNA replication, explaining how at later times the epigenome and cell fate can be altered.

**Highlights:**

1. The histone chaperone CAF-1 sustains DNA replication speed in single cells.
2. CAF-1 loss alters histone repertoire and delays chromatin maturation.
3. H3K9me3 and H3K27me3 regions respond differently to acute CAF-1 depletion.
4. Impaired S-phase chromatin assembly triggers an immediate response and a G0 arrest.

## INTRODUCTION

During every cell cycle, the entire genome must be accurately replicated. This process occurs in the context of chromatin, a highly organized DNA-protein complex that encodes epigenetic information essential for cell identity and survival^1^. Chromatin is organized in repetitive structural units called nucleosomes, which are composed of two copies of the histones H2A, H2B, H3, and H4 wrapped by 150bp of DNA. Histones and their different variants provide a platform for post-translational modifications (PTMs) that locally control chromatin organization and DNA accessibility^2^. Since these features regulate gene transcription and affect cell identity, faithful inheritance of chromatin during cell division is essential to safeguard the fate of the daughter cells^3–5^.

The inheritance of chromatin is tightly coupled to DNA replication during S-phase of the cell cycle^3–8^. Histone chaperones control chromatin dynamics in coordination with the replisome^5,9,10^. As DNA is copied, parental nucleosomes are recycled onto the two nascent DNA daughter strands to maintain local epigenetic information. In parallel, *de novo* chromatin assembly preserves chromatin density through the deposition of newly synthesized histones onto nascent DNA. These new histones initially lack local epigenetic PTMs, which are acquired after DNA replication through a process called chromatin maturation. Chromatin maturation mechanisms differ along chromosomes depending on the local environment ^11–14^, with heterochromatic histone PTMs accumulating with slower kinetics than the ones associated with active transcription^12^. This is despite heterochromatin-related factors being recruited rapidly on replicated DNA^14^. Moreover, to enable the rapid supply of chromatin components, S-phase cells coordinate the production of histone proteins with DNA replication^6,9,10,15,16^. Defects in chromatin replication are sensed by the cell cycle and affect cell proliferation^17–19^. Thus, S-phase chromatin dynamics safeguard epigenome stability and cell fate by orchestrating a complex network of diverse processes.

*De novo* chromatin assembly during DNA replication is a central part of this network^1,6–8,20,21^. This process is coordinated by the histone chaperone Chromatin Assembly Factor 1 (CAF-1)^22^. CAF-1 is a heterotrimeric complex (composed of CHAF1A, CHAF1B, and RBBP4) that binds newly synthesized H3-H4 and deposits them onto both replicated DNA strands^22–27^. CAF-1 is recruited to replication forks by the essential replication clamp PCNA^28–35^. Previous studies using knock-out or RNAi-mediated depletion of CAF-1 subunits have demonstrated a function for CAF-1 in safeguarding cell identity, epigenome stability, and DNA replication^20,33,36–41^. However, these depletion methods span multiple cell cycles, limiting our ability to understand the immediate mechanisms that drive these phenotypes. Therefore, it remains unclear how *de novo* chromatin assembly is mechanistically linked to genome replication and epigenome stability.

To understand these mechanisms, we need to study this process with higher temporal control. Highly controllable degron-systems allow the manipulation of essential proteins on a minute-to-hours’ timescale in cells^42,43^. Combined with single-cell genomics, quantitative proteomics and live-imaging technologies, these enable us to monitor immediate changes in chromatin organization, DNA replication and cell cycle progression, revealing the early mechanisms at the basis of these processes.

In this study, we find that acute CAF-1 degradation slows down the speed of DNA replication forks. This triggers a response that lowers histone mRNA levels and prolongs S-phase progression, without activation of canonical stress signals. These early events lead to changes in the availability of histone H3 variants and alter chromatin accessibility genome-wide with specific effects on facultative heterochromatin, which becomes transcriptionally de-repressed. After one S-phase in the absence of CAF-1, chromatin maturation is delayed and cells engage a p53 response that causes a cell cycle arrest in a G0 state after cell division. Our work uncovers the immediate mechanisms that are wired with *de novo* chromatin assembly during DNA replication and shows that CAF-1 safeguards epigenome stability within a single S-phase. The data suggest that rapid adaptations are required for cells to survive defects in this pathway over multiple cell divisions, as in cell fate trajectories.

## RESULTS

### Acute depletion of CAF-1 curtails DNA replication speed in single cells

To investigate the effects of acute depletion of CAF-1, we generated human RPE-1 cell lines with a bi-allelic endogenous knock-in of an N-terminal FKBP12^F36V^ tag^42^ on the largest subunit of CAF-1, named CHAF1A/p150 (referred to as degron-CHAF1A). This system enables rapid degradation of CHAF1A as early as 10 minutes after addition of the dTAG^V^-1 ligand (referred to as dTAG, Figure 1A and Supplemental Figure S1A). Since the CHAF1A subunit directly controls interactions with histones, DNA replication proteins and epigenetic factors^24,30,44–47^, our newly generated cell lines enable us to acutely inactivate CAF-1 and its key functionalities with high temporal resolution.

**Figure 1.**
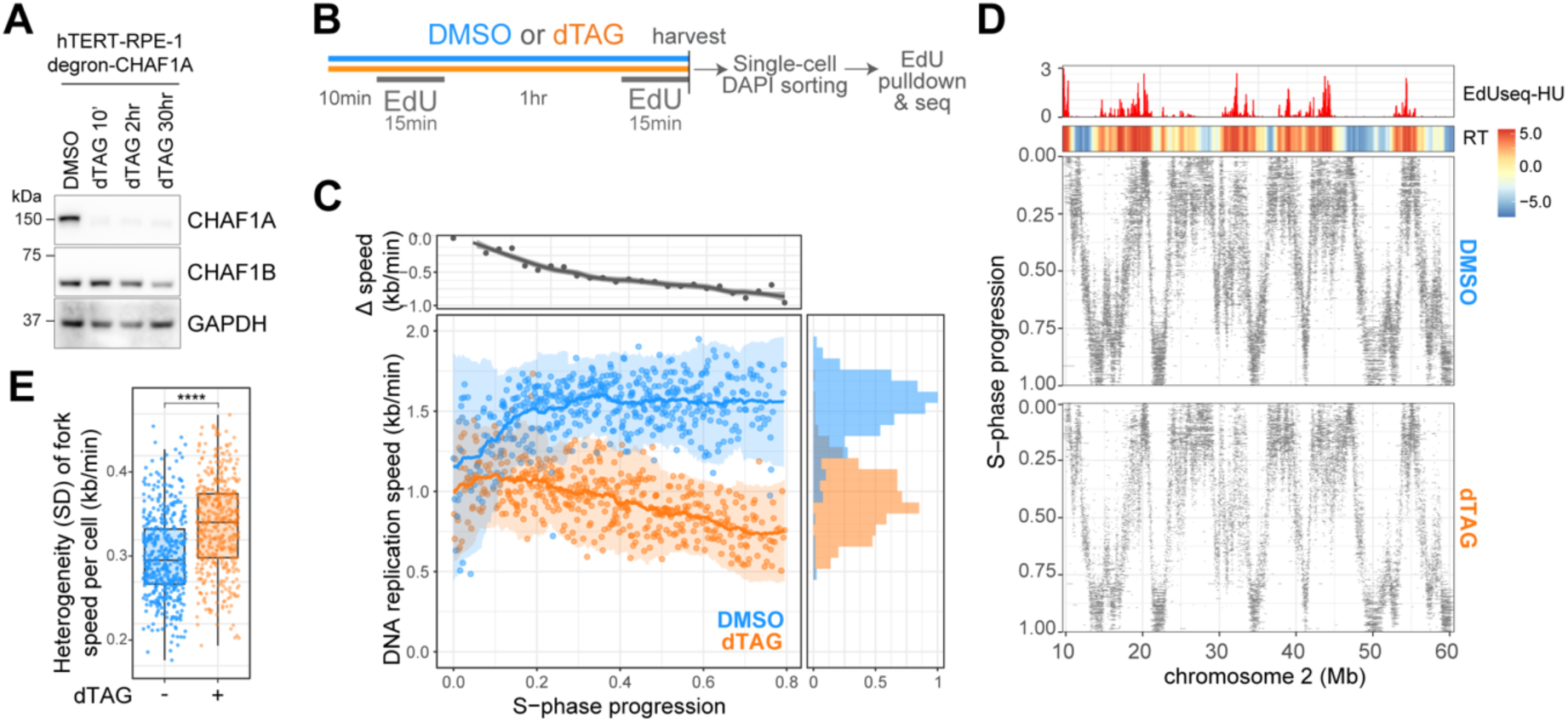
CAF-1 sustains DNA replication speed in single cells. A) Western blot analysis of RIPA extracted hTERT RPE-1 degron-CHAF1A cells treated with DMSO or dTAG for indicated time periods. B) Schematic representation of the treatment and sample preparation for scEdU-seq experiments. C) DNA replication speed over S-phase in degron-CHAF1A treated with DMSO (blue) or dTAG (orange) subjected to EdU labeling scheme (see Figure 1B). Difference in DNA replication speeds between DMSO and dTAG in kb/min (y-axis) over S-phase progression (x-axis, top-left), marginal normalized density (x-axis) of DNA replication speed in kb/min (y-axis) colored for DMSO- (blue) or dTAG-treated cells (orange, bottom-right). Each dot represents a single cell. D) Heatmap showing scEdU-seq maximum normalized log counts for DMSO-treated and dTAG-treated RPE-1 degron-CHAF1A cells ordered according to S-phase progression (y-axis) and binned per 40 kb bins (x-axis) for a 50 megabase region of chromosome 2. Top: heatmap showing log2-fold ratio of early to late Repli-Seq indicating replication timing (top) and bar graph showing EdU-seqHU, to mark replication origins, of the same stretch of chromosome 2 (bottom). E) Boxplots showing heterogeneity of fork speed within single cells expressed as the standard deviation of DNA replication speeds (kb/min, y-axis) from degron-CHAF1A cells treated with DMSO or dTAG (x-axis). Adjusted p-values were determined by Pairwise T-test and corrected Bonferroni multiple testing correction.

CAF-1 acts during DNA replication and its depletion affects EdU incorporation levels (Supplemental Figure S1B) as previously observed^21,30,48^, indicating that it can influence DNA synthesis. Therefore, we first set out to determine how DNA replication is affected by acute depletion of CAF-1. To this end, we applied scEdU-seq on degron-CHAF1A cells to monitor location and speed of replication forks genome-wide with single-cell resolution^49^. The experiment is carried out on asynchronous cells with acute depletion of CAF-1 for just 10 minutes prior to EdU labeling (Figure 1B). The single-cell resolution provides quantitative information on DNA replication both temporally along S-phase, as well as spatially on the genome, removing bias from averaging effects.

We find that CAF-1 depletion leads to a persistent decrease in replication speed genome-wide during S-phase progression (Figure 1C). This effect is strongest in late S-phase, when heterochromatic regions are replicated^50^ (Figure 1C). Interestingly, there are no global alterations in DNA replication timing (i.e. changes in early, mid and late replicating regions) upon CAF-1 loss (Figure 1D and Supplemental Figure S1C-D). Moreover, we measure a similar number of replication forks per cell (Supplemental Figure S1E-F) and a similar percentage of cells displayed robust DNA replication profiles across S-phase (Supplemental Figure S1G). Based on these observations, we conclude that acute depletion of CAF-1 slows down the speed of replication forks, without halting DNA replication. Interestingly, we also observe an increase in the standard deviation of the speed of replication forks within single cells (Figure 1E), indicating that fork speed becomes more heterogeneous, with a stronger effect in late S-phase cells (Supplemental Figure S1H). To our surprise, the effect of CAF-1 loss on replication speed is not exacerbated over time, as removal of CAF-1 for longer periods (two hours prior to EdU labeling) results in a comparable slowdown (Supplemental Figure S1I) with no effects on replication timing (Supplemental Figure S1J). Taken together, these data demonstrate that CAF-1 sustains DNA replication speed.

### Acute CAF-1 depletion prolongs S-phase without triggering ATM or ATR activation

Next, we asked how the slowdown of replication forks affected S-phase progression. To this end, we monitored cell cycle progression by live-microscopy at a single-cell level using a fluorescent fragment of DNA helicase B (DHB) as CDK2 activity sensor^48^. CDK2 activity builds up during interphase and quickly drops at the onset of mitosis, allowing us to monitor the duration of these events along the cell cycle in asynchronously cycling cells^48^. By aligning single cells according to the time of dTAG or DMSO addition relative to their last anaphase, we evaluated the effect of CAF-1 loss dependent on the different cell cycle phases. Strikingly, acute CAF-1 loss in early or late S-phase causes an immediate plateau in CDK2 activity (Figure 2A-B and Supplemental Figure S2A-B). Conversely, cells that lose CAF-1 in G1 phase do not respond immediately, they ramp up CDK2 activity normally until they enter S-phase (Figure 2C and Supplemental Figure S2C). Moreover, cells where CAF-1 is depleted in G2 do not change CDK2 activity until their daughter cells reach the next S-phase (Figure 2D and Supplemental Figure S2D). These data demonstrate that CAF-1 depletion is rapidly sensed only during S-phase, where it leads to a dampening of CDK2 activity build-up. Importantly, in line with DNA replication not stopping, we do not observe an S-phase arrest. The dampening of CDK2 activity is sustained until entry into mitosis, which we observe in all cells lacking CAF-1 (Supplemental Figures S2A-D). Flow cytometry analysis confirmed these observations showing that CAF-1-depleted cells spend more time in S-phase after release from CDK4/6 inhibition, used to synchronize the cells in G1^51^ (Supplemental Figure S2E). Thus, CAF-1 loss slows down DNA replication speed, thereby prolonging S-phase. This results in an increase in cell cycle duration from 20 to 30 hours in control and dTAG conditions, respectively (Figure 2A-D).

**Figure 2.**
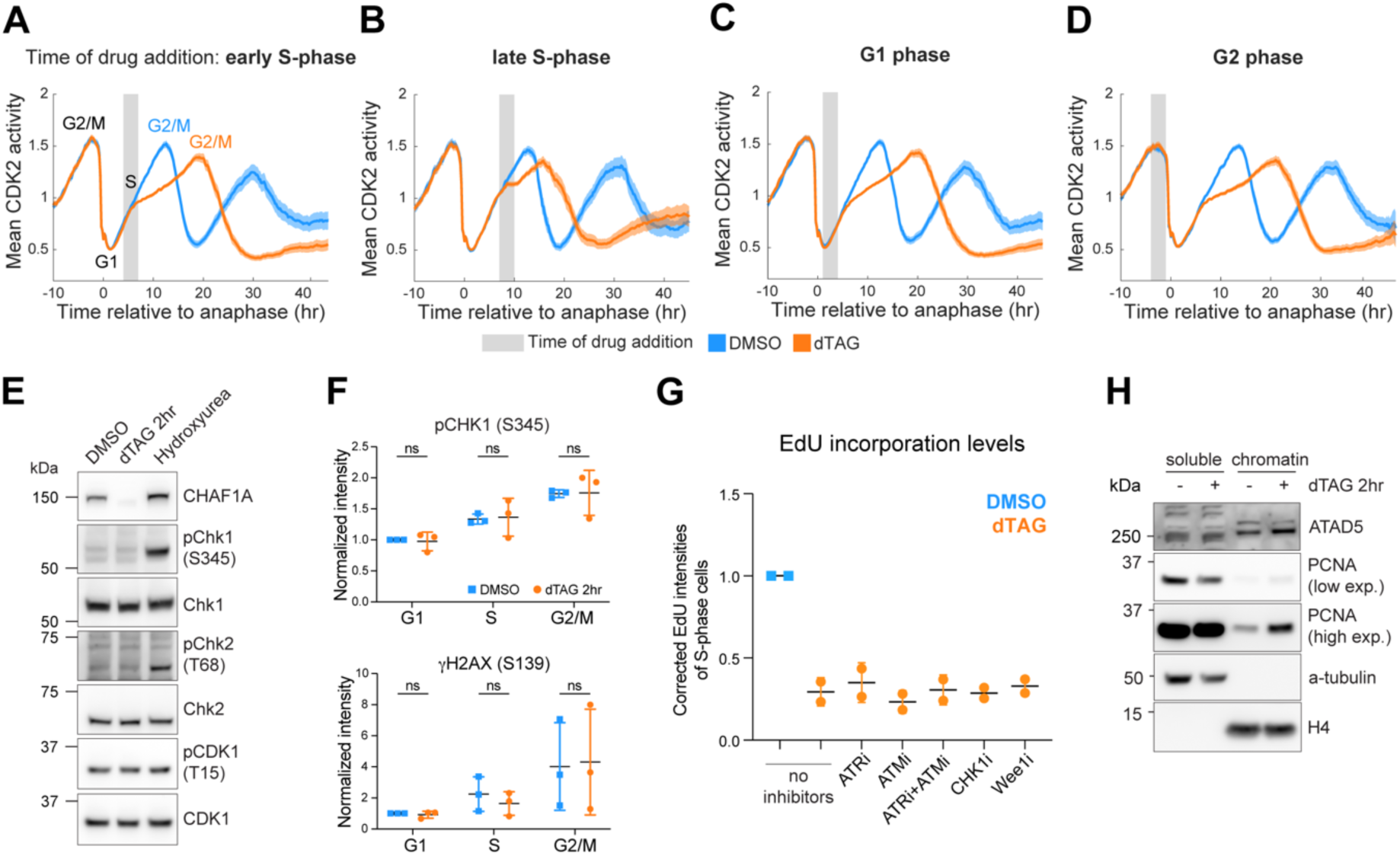
Acute CAF-1 depletion prolongs S-phase without triggering ATM or ATR activation. A-D) Mean signal of CDK2 activity sensor aligned computationally to the time of anaphase. Cells included in the different analyses experienced the start of DMSO (blue) or dTAG (orange, 1uM) treatment relative to their last anaphase in indicated cell cycle phases. E) Western blot analysis of RIPA extracted degron-CHAF1A cells treated for 2hr with DMSO, dTAG or HU=Hydroxyurea (10mM) probed with indicated antibodies. F) Flow cytometry analysis for pChk1 (S345) and ɣH2AX (S139) intensities (y-axis) for indicated cell cycle phases (x-axis) in RPE-1 degron-CHAF1A cells treated for 2hr with DMSO (blue) or dTAG (orange). Error bars represent SD as result of unpaired, parametric t-tests (ns=non-significant). G) Corrected EdU intensities (y-axis) for DMSO or dTAG (2hr) degron-CHAF1A S-phase cells treated with indicated inhibitors (x-axis). H) Western blot of soluble and chromatin fraction of RPE-1 degron-CHAF1A cells treated with DMSO or dTAG (2hr) probed with indicated antibodies.

To investigate the mechanism of DNA replication slowdown, we tested whether cell cycle checkpoints are activated upon acute CAF-1 depletion. Prior studies using long-term (i.e., 48-72hr) depletion or mutations of CAF-1 in flies, human or mouse cells implied checkpoint activation^18,52–54^, while others reported no effects on either DNA damage checkpoints or cell cycle progression^21,48,55^. Acute depletion of CAF-1 in our experimental setting did not cause an increase in phosphorylation of CHK1 (Ser345), CHK2 (Thr68), H2AX (Ser139), CDK1(Tyr15) or RPA (Ser4/8) (Figure 2E-F and Supplemental Figure S2F). Moreover, inhibition of the DNA damage kinases ATR or ATM, as well as Wee1 kinase, was unable to reverse the reduction of EdU intensity seen upon CAF-1 depletion (Figure 2G). These data indicate that acute depletion of CAF-1 does not trigger an ATM or ATR response.

However, we did observe an accumulation of PCNA on chromatin upon acute CAF-1 loss, without significant depletion of the available soluble PCNA pool (Figure 2H). Interestingly, we also observed increased chromatin levels of the PCNA unloading complex ATAD5 (Figure 2H), indicating that PCNA accumulation is not due to unavailability of the unloader. Thus, PCNA dynamics are rapidly altered upon CAF-1 degradation, similar to what was previously observed upon perturbation of other players in chromatin dynamics during DNA replication^21,56,57^. We show PCNA accumulates as early as 2 hours following CAF-1 degradation, and this also involves ATAD5, suggesting that PCNA may have a direct function in the slowdown of replication forks.

### Transcriptional response of S-phase cells to CAF-1 depletion

To understand the immediate cellular response to acute CAF-1 depletion, we sought to analyze the transcriptional effects upon defective *de novo* chromatin assembly in correlation to the different cell cycle phases. To this end, we performed single-cell RNA sequencing using VASA-seq, which enables sequencing of all RNA, including non-polyadenylated genes (e.g. histone genes) throughout the cell cycle^58^. This allowed us to analyze changes in the transcriptional profile of short-term CAF-1 depleted cells (2hr) per cell cycle phase. In addition to control treatment with DMSO, we also used long-term CAF-1 depleted cells (30hr), which are expected to display a strong and diverse transcriptional response^36,44^. To analyze the VASA-seq data, we used Leiden clustering to uncover cell clusters (Figure 3A and Supplemental Figure S3A), followed by a Uniform Manifold Approximation and Projection (UMAP) to visualize the cell cycle states per condition (Figure 3B and Supplemental Figure S3B-C)^58^. CAF-1 depleted cells behave distinctly from DMSO-treated and WT cells. Particularly, upon short-term CAF-1 depletion (2hr), only S-phase cells are affected, as evidenced by their separation from DMSO-treated cells (gray dotted line, compare clusters 3 and 7) (Figure 3A and 3C and Supplemental Figure S3A). This confirms that CAF-1 loss is rapidly sensed in S-phase and alters cell cycle states. Upon longer CAF-1 depletion (30hr), more cell cycle phases are affected, with the rising of unique G0/G1 and S/G2 populations (see clusters 6 and 9, respectively) (Figure 3A and D and Supplemental Figure S3A-C). These analyses support the notion that inactivation of CAF-1 leads to immediate transcriptional effects only during S-phase. Conversely, prolonged CAF-1 loss more broadly affects the cell cycle. Thus, CAF-1 is closely wired into S-phase regulatory mechanisms, in line with its role in *de novo* chromatin assembly during DNA replication ^25,28,52^.

**Figure 3.**
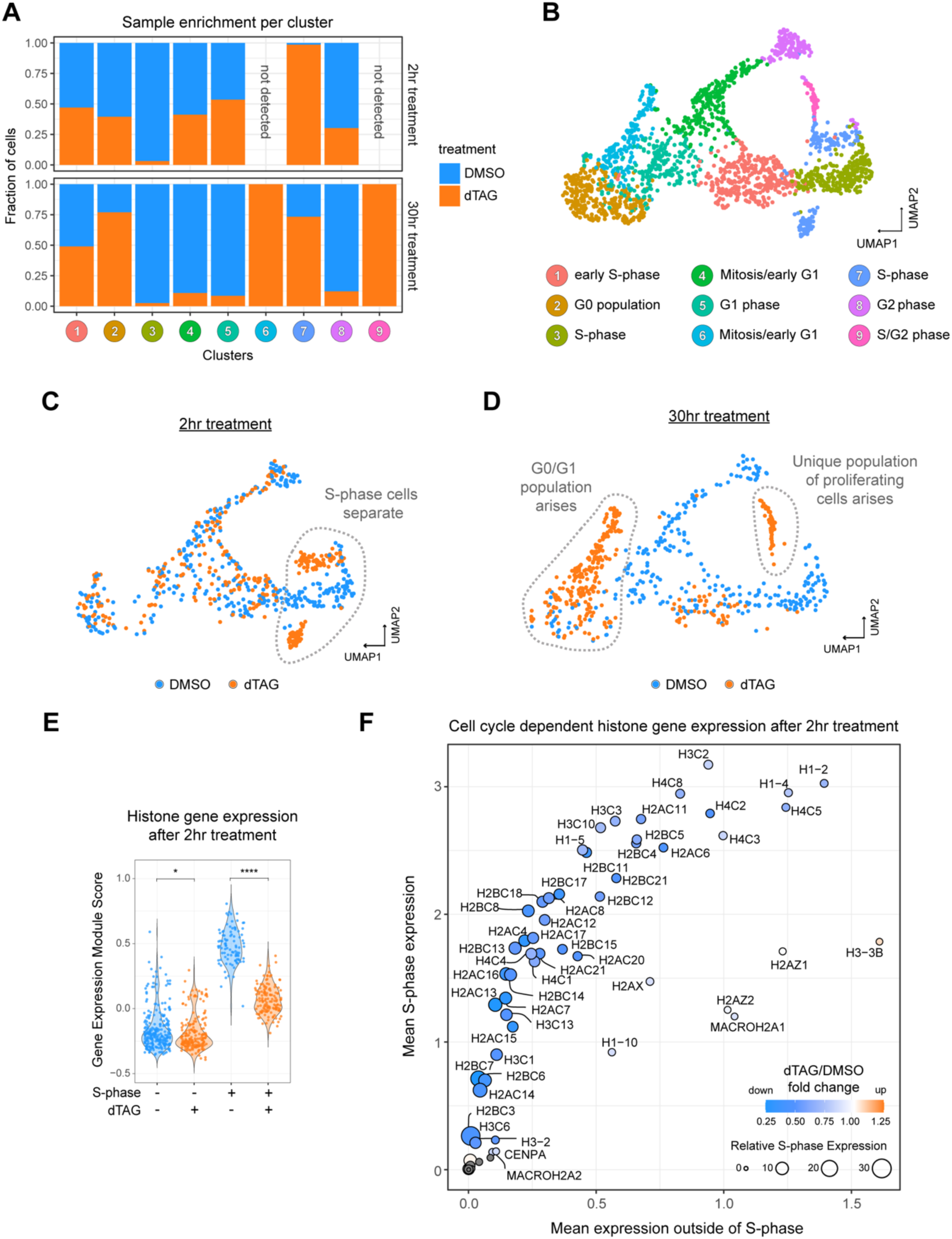
CAF-1 controls the levels of histone mRNA in S-phas. A) Stacked bargraph displaying the normalized contribution of each experimental condition to each Leiden cluster (x-axis). B) UMAP plot obtained from total gene-wise RNA counts of sequenced cells analyzed with Seurat, colored by cycle stages identified by unsupervised clustering and characterized with marker genes for cell cycle progression shown in Figure S3B. Each dot represents a single cell. C-D) UMAPs determined using total RNA counts per gene illustrating cells treated for 2hr (C) or 30hr (D) with DMSO or dTAG. Differences are highlighted in gray. E) Violin plot of the single-cell expression analysis for all histone genes using ModuleScore function from Seurat for 2hr DMSO or dTAG treated degron-CHAF1A cells split by non S-phase and S-phase clusters. Adjusted p-values were obtained by Pairwise T-test followed by Bonferroni multiple testing correction. See Table S1. F) Analysis of CAF-1 dependent expression of all histone genes. In this scatterplot, each histone gene is represented by a dot, which is colored by the fold-change in expression in acute CAF-1 depletion (2hr, dTAG/DMSO). Gray denotes genes not detected in DMSO or dTAG samples. Their size indicates the relative S-phase expression (S-phase/outside of S-phase expression) defined as a fold enrichment per histone gene. The x-axis represents the mean expression of each histone gene outside of S-phase and the y-axis represents the mean expression of each histone gene in S-phase.

### CAF-1 controls the levels of histone mRNA in S-phase

Since acute CAF-1 depletion specifically affects S-phase cells, we investigated the top 20 differentially expressed genes in this condition and observed a dramatic downregulation of histone mRNAs (Supplemental Figure S3D). Indeed, global expression of all histone genes significantly decreased in single S-phase cells (Figures 3E, Supplemental Figure 3E, Table S1). We validated this finding by measuring lowered expression of histone H4.2 using RNA FISH in S-phase cells (Supplemental Figure S3F). When further analyzing the expression levels of all histone genes, we found that this phenotype was dominated by a specific downregulation of the mRNA of replicative histones, such as H3.1/2 and H4 (Figure 3F and Supplemental Figure S3G), whose expression is mostly restricted to S-phase. In sharp contrast, mRNA levels of non-replicative histone variants, such as H3.3, macroH2A, and H2AZ did not decrease upon depletion of CAF-1 (Figure 3F and Supplemental Figure S3H). In fact, we observed that H3.3 is mildly upregulated in CAF-1 depleted cells. This indicates that CAF-1 depletion affects the levels of replicative histone mRNAs. These data show that replicative histone mRNA levels in S-phase are linked to CAF-1 function, similar to previous observations focusing on players of the new histone supply pathway^8,21,53,59–61^. Interestingly, in those studies, 2-day RNAi depletion of CHAF1B did not affect histone mRNAs^59^. This may indicate that CHAF1A and CHAF1B have different functions within S-phase. Another plausible explanation may arise from the distinct timing of the experiments and the single-cell resolution of our approach. As CHAF1A is the functionally defining subunit of the CAF-1 complex ^24,30,44–47^, our findings argue that CHAF1B depletion may not fully recapitulate CAF-1 loss.

### CAF-1 loss leads to histone imbalance and delays chromatin maturation within a single S-phase

To directly measure how the changes in histone mRNAs upon CAF-1 loss affect histone composition and histone PTMs in chromatin, we used a SILAC-based quantitative mass spectrometry approach, which measures newly synthesized (i.e. new) and parental (i.e. old) H3 and H4 variants and their PTMs (Figure 4A and Supplemental Figure S4A-B)^62^. We quantified the relative abundance of histone H3 and H4, in addition to the replicative H3.1/2 and non-replicative H3.3 histone variants after a single S-phase without CAF-1 (Figure 4B). In line with a reduction in histone mRNAs, we found that in the absence of CAF-1, there is a decrease in newly synthesized histone H3.1 (Figure 4B). Conversely, we find an increase in newly synthesized H3.3 protein (Figure 4B), as assessed by quantifying H3.3-specific peptides. The increase in H3.3 protein was confirmed by western blot (Figure 4C). As 98% of cellular histones are incorporated into chromatin^63^, our data suggests that chromatin is enriched in H3.3 upon CAF-1 loss, which likely compensates for the decrease in canonical H3.1/2 histones. This is in line with previous observations that RNAi-depletion of CAF-1 increases global H3.3 incorporation^64,65^. Notably, we show that this is a rapid cellular response, as it occurs as early as the first S-phase without CAF-1.

**Figure 4.**
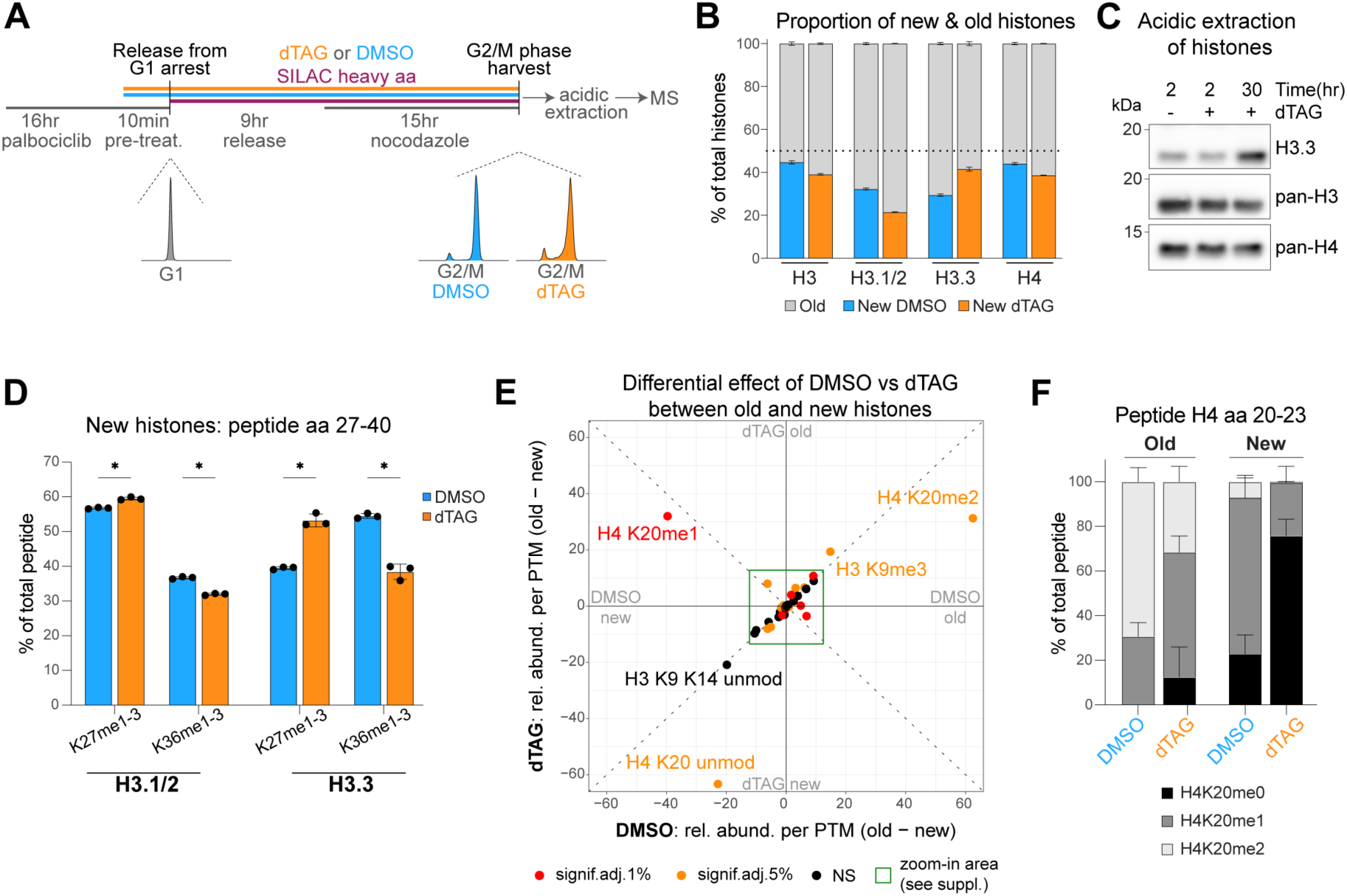
CAF-1 loss leads to histone imbalance and globally affects chromatin maturation within a single S-phase. A) SILAC-MS set up for analysis of old and new histones, their PTMs and variants of cells that underwent exactly one full S-phase after the start of DMSO/dTAG treatment. DAPI FACS profiles of the different conditions are shown. B) Cumulative barplots of the proportion of old and new histones in DMSO and dTAG. The horizontal dotted line is set at 50%. C) Western blot of acidic extractions of degron-CHAF1A cells (clone 8) treated with DMSO or dTAG for indicated times. D) Quantification of new H3.1/2 and H3.3 peptides (amino acids 27-40). The height of the barplots represent the relative abundances of mono-, di- and tri-methylation of K27 or K36 that were summed up. E) Scatterplot of the differential effect of DMSO (x-axis) vs dTAG (y-axis) between old and new histones. Colors encode significance levels (p-value) after Benjamini-Hochberg adjustment. A zoom-in is available in Supplemental Figure S4D. F) Quantification of the H4K20-anchored PTMs relative abundance on old and new H4 peptide 20-23 associated with DMSO or dTAG treatment. (B,D,F) Quantification of histones and their PTMs of degron-CHAF1A cells (clone 8) treated as described in (A). Percentages are relative to the total intensity of related peptides and averaged across n = 3 biological replicates. Error bars represent SD, asterisks represent p-values as * < 0.05, ** < 0.01, *** < 0.001, **** < 0.0001 as result of unpaired, parametric t-tests.

Furthermore, we analyzed histone PTMs on the unique peptide that distinguishes the tail of H3.1/2 or H3.3. This peptide encompasses residues K27 and K36, which are established targets of epigenetic marks^66^, where H3K27 is methylated in transcriptionally silenced regions and H3K36 in active domains. We found significant changes in the relative abundance of H3K27 and H3K36 methylation status in old and new H3.1/2 and H3.3 (Figure 4D and Supplemental Figure S4C). However, the most profound effect was seen on newly synthesized H3.3 peptides. In control DMSO conditions, new H3.3 shows a higher percentage of H3K36 methylation compared to H3K27 methylation, likely given by its rapid incorporation in transcriptionally active regions during S-phase. In the absence of CAF-1, new H3.3 peptides reversed this distribution displaying higher percentage of H3K27 methylation (Figure 4D). Interestingly, this dTAG-dependent distribution of H3K27 and H3K36 methylation on H3.3 closely resembles the distribution of these marks on H3.1/2 (Figure 4D). Thus, upon CAF-1 depletion newly synthesized H3.3 mimics the H3.1/2 distribution of H3K27 and H3K36 methylation status, implying that in cells lacking CAF-1, newly synthesized H3.3 is incorporated in genomic regions where, in unperturbed conditions, H3.1/2 resides.

Next, we analyzed global changes in other histone H3 and H4 PTMs. Several histone PTMs are significantly affected by CAF-1 depletion, most notably H3K4me1, H3K18ac, acetylation of the H4 N-terminal tail (aa 4-17) and the heterochromatic mark H3K9me3 (Figure 4E and Supplemental Figure S4D-E, Table S2). However, the most pronounced effects were observed for the methylation status of H4K20 (Figure 4E-F). H4K20 methylation functions as a timer for histone age, from their synthesis in early S-phase (H4K20me0) to their maturation on chromatin later starting from late G2 (H4K20me1/2)^12,67–70^. We found a relative increase in H4K20me0 and a corresponding relative decrease in H4K20me2, both on new and old histones (Figure 4F). This suggests that chromatin is “younger” (i.e. less mature) after a single S-phase in the absence of CAF-1. Interestingly, its effect involves not only the pool of newly synthesized histones, but also the parental (i.e. old) histones in chromatin, indicating that *de novo* chromatin assembly during S-phase has the potential to alter epigenetic states within a single S-phase.

Together, these data show that within a single S-phase CAF-1 loss alters histone protein levels with consequences for global chromatin composition at the end of S-phase.

### CAF-1 assembles nascent chromatin genome-wide

To understand the mechanisms that lead to these global changes in chromatin composition, we set out to monitor how chromatin organization along the genome is affected by CAF-1 depletion during S-phase. To this end, we applied repli-ATAC-seq, which combines Tn5-mediated tagmentation of accessible DNA with EdU incorporation to measure chromatin accessibility specifically at replicated regions^71^. In combination with acute CAF-1 depletion, this approach allows us to directly measure the role of CAF-1 in assembling replicated chromatin throughout the genome with high temporal resolution. Using this technique, we analyzed 1) nascent chromatin to observe CAF-1-dependent effects during DNA replication, 2) mature chromatin to measure the effects of prolonged CAF-1 loss during and after DNA replication and 3) post-replication chromatin to observe the role of CAF-1 after DNA replication (Figure 5A and Supplemental Figure S5A). All samples were mixed with EdU-labeled S2 *Drosophila* cells as spike-in to allow normalization and comparison of accessibility changes between conditions. These spike-in were also used to exclude that the lower EdU incorporation rates upon CAF-1 depletion (Supplemental Figure S1B) affected the repli-ATAC-seq analyses (Supplemental Figure S5B).

**Figure 5.**
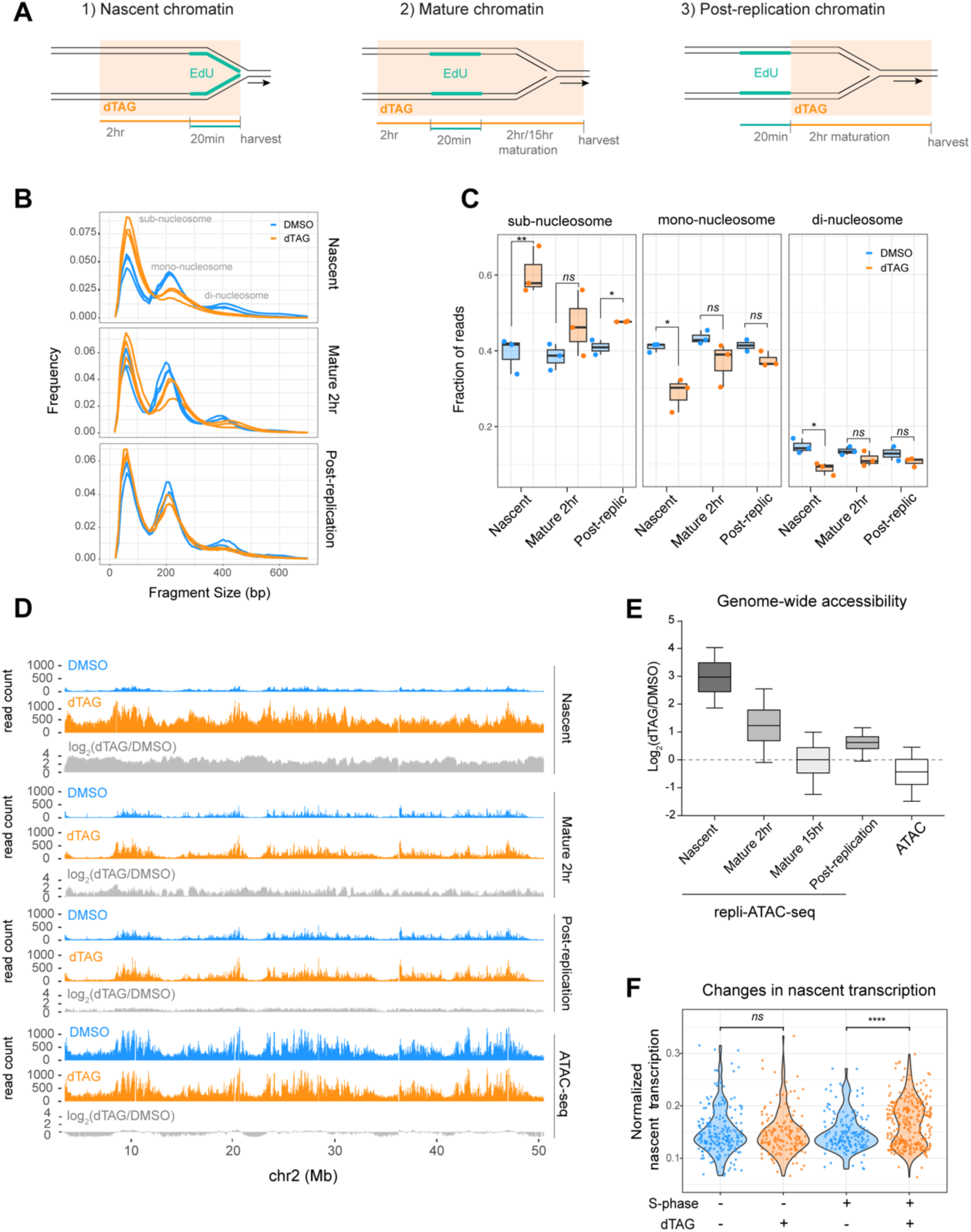
CAF-1 controls chromatin assembly and transcriptional fidelity in S-phase. A) Schematics of repli-ATAC-seq conditions. B) Fragment size frequency of repli-ATAC-seq samples for nascent, mature 2hr and post-replication chromatin from three independent experiments. C) Boxplot quantification of reads within the sub-nucleosomal (<150 bp), mono-nucleosomal (150-300 bp) and di-nucleosomal (300-450 bp) size range relative to the total amount of DNA fragments. P-values were calculated using binomial-test. D) Spike-in normalized signal (y-axis) of nascent, mature and post-replication repli-ATAC-seq samples and ATAC-seq sample over a selected region of about 50 Mb on chromosome 2. In blue DMSO control samples, in orange dTAG treated samples, in grey Log2 fold-change quantification of dTAG over DMSO signal. E) Boxplot quantifying accessibility signal changes of dTAG over DMSO over the whole genome (15 kb binned genome) in different repli-ATAC-seq conditions and ATAC-seq (2hr dTAG treatment). F) Violin plots of single cell nascent transcription quantification as determined by the number of unique spliced (mature) total RNA reads divided by the number of unique unspliced (nascent) total RNA (y-axis) as produced by the RNA Velocity algorithm for CHAF1A cells treated for 2 hours with DMSO and dTAG split by non S-phase and S-phase clusters (x-axis). Adjusted p-values were obtained by Pairwise T-test followed by Bonferroni multiple testing correction.

In nascent chromatin, CAF-1 depletion leads to an increase in sub-nucleosomal fragments and to a decrease in mono- and di-nucleosomal fragments (Figure 5B-C). This strongly suggests a loss of nucleosomes on nascent DNA. In contrast, in mature or post-replication chromatin, CAF-1 depletion has milder effects, mainly leading to an increase in linker length between nucleosomes (i.e. mono- and di-nucleosomal peaks shift to higher sizes) (Figure 5B-C). These data show that CAF-1 significantly alters chromatin on nascent DNA, as expected^24,48^.

To quantify genome-wide effects of CAF-1 on chromatin accessibility, we monitored the fold-change of the repli-ATAC signal in dTAG over DMSO samples (log_2_(dTAG/DMSO)) per 15Kb-bin and plotted its distribution. In nascent chromatin, 99% of the genome displays a positive fold-change enrichment (Figure 5D-E and Supplemental Figure S5C), demonstrating that acute CAF-1 depletion results in a dramatic gain of accessibility on newly replicated DNA throughout the genome. This effect decreases over time after DNA replication, with an intermediate increase in accessibility after 2 hours of chromatin maturation (Mature 2hr, Figure 5E), and a near-complete recovery of accessibility after 15 hours of chromatin maturation (Mature 15hr, Figure 5E and Supplemental Figure S5C). These data indicate that during chromatin maturation, CAF-1 independent backup mechanisms facilitate nucleosome assembly, as previously proposed^64^. Post-replication removal of CAF-1 results in a limited increase in genome-wide chromatin accessibility (Figure 5D-E), confirming that the main role of CAF-1 is during DNA replication. Moreover, we performed ATAC-seq after a 2-hour depletion of CAF-1 to assess the chromatin accessibility changes at steady-state level and detected no global alterations (Figure 5D-E). Similar results were obtained when analyzing the parental non-replicated regions from repli-ATAC-seq samples, which also showed no global alteration in chromatin accessibility (Supplemental Figure S5C-D). These data demonstrate that CAF-1 controls chromatin assembly during DNA replication at a genome-wide scale, with no major roles in nucleosome assembly outside of DNA replication.

### CAF-1 controls transcriptional fidelity in S-phase

Next, we investigated if the opening of nascent chromatin upon acute CAF-1 loss alters nascent transcription. To this end, we analyzed the ratio of unspliced over spliced RNA in our scRNA-seq data after short-term CAF-1 depletion. We found an increase in nascent transcription specifically in S-phase cells (Figure 5F). Thus, defective *de novo* chromatin assembly during DNA replication leads to loss of transcriptional fidelity during S-phase, suggesting that proper chromatin assembly affects the transcriptional state of genes behind replication forks. To see if this was due to changes in the accessibility of annotated promoters, genic and intergenic (e.g., enhancers) regions, we analyzed our repli-ATAC-seq datasets and found that accessibility of these regions is altered in nascent chromatin, in line with the global accessibility changes observed earlier (Supplemental Figure S5E). This supports the notion that the opening of nascent chromatin observed upon CAF-1 depletion promotes a loss of transcriptional fidelity, as also seen recently in *Saccharomyces cerevisiae* CAF-1 knock-out strains^72^.

Taken together, CAF-1 acts globally to assemble chromatin during DNA replication across the entire genome. Via this function, CAF-1 safeguards transcriptional fidelity of the replicated genome. These data indicate that back-up nucleosome assembly pathways may promote chromatin compaction, but that they are not sufficient to restore functional chromatin organization. This is also supported by the global changes we observe in histone variants and PTMs after a single S-phase without CAF-1 (Figure 4).

### Acute CAF-1 loss differentially affects constitutive and facultative heterochromatin

Previous CAF-1 RNAi or knockout studies have shown loss of heterochromatin stability after multi-day treatments^33,44,48,73–76^. In our proteomic data, CAF-1 loss has a significant effect on the constitutive heterochromatin H3K9me3 modification, where this mark increases in old histones and decreases in new ones upon CAF-1 depletion (Figure 4E and Supplemental Figure S4E). Moreover, the facultative Polycomb-associated H3K27me3 mark significantly increases on all histones with strongest effects on the new H3.3 variant (Figure 4D and Supplemental Figure S4C-D). To understand how CAF-1 contributes to these changes and the reported phenotypes, we monitored chromatin accessibility within these regions in our different repli-ATAC-seq datasets. In nascent chromatin, both heterochromatic regions showed a significant increase in accessibility compared to the rest of the genome during DNA replication (Figure 6A-B and Supplemental Figure S6A). However, the CAF-1 dependent opening of H3K9me3-marked regions is more dramatic than in H3K27me3 domains (Figure 6A-B and Supplemental Figure S6A, nascent samples). These data indicate that the initial compaction of both heterochromatic regions is particularly susceptible to CAF-1 loss during DNA replication.

**Figure 6.**
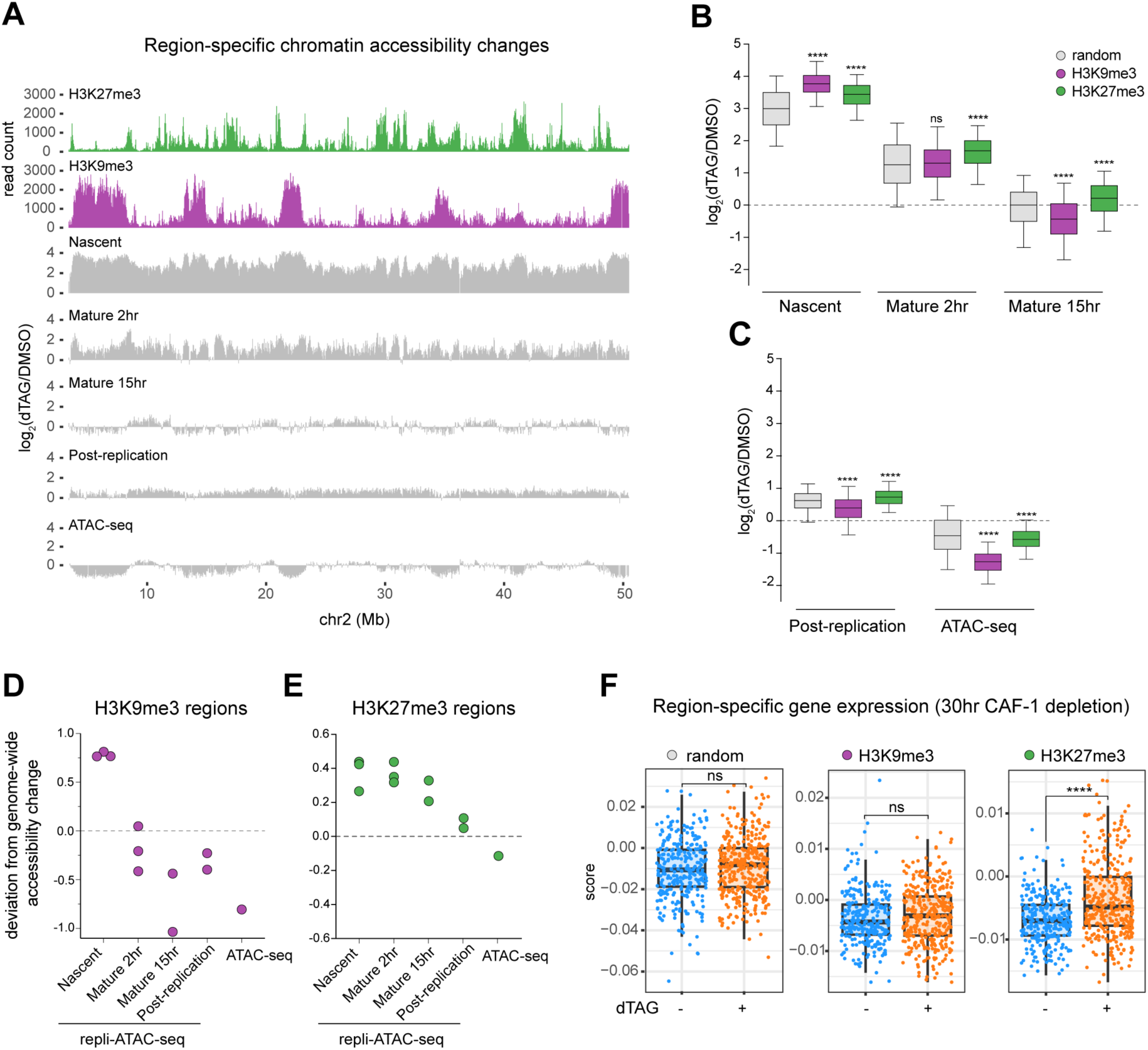
Acute CAF-1 loss differentially affects constitutive and facultative heterochromatin. A) Genomic tracks of ChIC-seq signal for H3K9me3 (purple) and H3K27me3 (green) in DMSO treated cells. In grey: tracks showing changes in accessibility following CAF1 depletion for repli-ATAC and ATAC-seq experiments, over 50 Mb loci on chromosome 2. B-C) Boxplots of changes in accessibility signal, upon CAF-1 removal, per region, in nascent, mature (2hr or 15hr), post-replication repli-ATAC-seq samples, and in ATAC-seq (2 hr dTAG). Each box plot represents 1000 bins either randomly selected genome-wide (grey) or selected by the highest ChIC-seq signal for H3K9me3 (purple) and H3K27me3 (green) regions. One-way ANOVA test was performed to determine statistically significant differences between each genomic regions within each sample. Correlation analysis over the whole genome is shown in Supplemental Figure S6A. D-E) Quantification of the differences between the median of the randomly selected bins and the median of the heterochromatic regions (H3K9me3 in purple, H3K27me3 in green), from panels B and C. Wilcoxon-Test was used to extract the median for each genomic region within sample and used to quantify its deviation from the genome-wide accessibility signal for H3K9me3 and H3K27me3 regions in all repli-ATAC-seq samples and ATAC-sample. F) Boxplots of single cell expression analysis using ModuleScore function from Seurat. Cells were treated with 30hr of DMSO (blue) or dTAG (orange). Panels show module expression levels for 250 random genes in the genome (left panel) or in 250 genes with highest accessibility scores in H3K9me3 regions (middle panel) and 250 genes with highest accessibility scores in H3K27me3 regions (right panel). See Table S1.

After DNA replication, constitutive H3K9me3 heterochromatin starts to compact during chromatin maturation (2hr), independently of CAF-1. This compaction stabilizes at longer chromatin maturation times (15hr), reaching the accessibility levels observed in the ATAC-seq sample (Figure 6A-C and Supplemental Figure S6A-C). These observations indicate that, while H3K9me3 heterochromatin opens up more dramatically during DNA replication without CAF-1, other mechanisms rapidly compensate for CAF-1 loss after DNA replication to promote its compaction. This compaction appears exacerbated by the absence of CAF-1, also at steady-state conditions (Figure 6D). In contrast, the less drastic chromatin accessibility defects in H3K27me3 domains observed in nascent chromatin persist during chromatin maturation (Figure 6E). Thus, CAF-1-independent chromatin maturation mechanisms are unable to rescue chromatin compaction defects at H3K27me3 domains, at least up to 15 hours after DNA replication. Strikingly, the opening of these regions is not observed outside of DNA replication, such as at steady-state conditions (i.e. ATAC-seq after 2hr dTAG treatment) or in the post-replication or unbound samples (Figure 6E). These controls indicate that the defects in compaction of H3K27me3 regions are strictly dependent on CAF-1 function during DNA replication.

Taken together, H3K9me3 and H3K27me3-marked heterochromatin are highly susceptible to CAF-1 loss during DNA replication, displaying increased chromatin accessibility defects on nascent DNA. Interestingly, compaction of H3K9me3 chromatin can be recovered post-replication by CAF-1-independent mechanisms, while H3K27me3 regions persist in a hyper-accessible state for over 15 hours after DNA replication. Moreover, CAF-1 loss may increase constitutive heterochromatin compaction at steady-state level, raising the possibility of local CAF-1 activities in these regions beyond DNA replication.

### Sustained CAF-1 depletion results in loss of transcriptional fidelity in facultative heterochromatin

To understand the functional consequences of these changes in chromatin accessibility, we monitored the distribution of the heterochromatic histone PTMs along the genome after CAF-1 loss. Neither H3K27me3 nor H3K9me3 displays significant changes after long-term (15 or 30hr) CAF-1 depletion, as seen in ChIC-seq^77^ (Supplemental Figure S6D-E). Thus, domain-level distributions of these histone marks do not change within a single S-phase. However, when we analyzed whether these regions are transcriptionally de-regulated in our scRNA-seq data, we found that transcripts originating from H3K27me3 regions are slightly, but significantly, increased after 30hr of CAF-1 depletion (Figure 6F and Table S1). This effect is specific to H3K27me3 domains, as a randomized geneset or transcripts from H3K9me3 regions do not show such an increase in transcript levels (Figure 6F and Table S1). These data show that sustained accessibility of H3K27me3 domains after DNA replication without CAF-1 can alter their transcriptional state, while CAF-1-indepedenent mechanisms safeguard the integrity of H3K9me3 regions, at least within the first 30 hours. Moreover, our data suggest that the loss of transcriptional fidelity may result from changes beyond histone PTM distribution at a domain-wide level. In fact, our mass-spectrometry experiments showed that newly synthesized H3.3 acquires H3K27me1-3 modifications upon CAF-1 loss (Figure 4D), suggesting the deposition of this histone variant within these domains. Thus, histone H3.3 variants incorporation could in part explain the functional changes in H3K27me3 regions.

Taken together, our data show that by assembling chromatin during DNA replication, CAF-1 safeguards chromatin compaction at replication forks. Within a few hours after DNA replication, heterochromatic regions become hyper-accessible upon CAF-1 loss. After DNA replication, constitutive and facultative heterochromatin differentially recover from this opening effect, likely due to locally acting CAF-1-independent chromatin maturation mechanisms^12–14^. The changes in histone repertoire that we uncovered with our proteomic analysis likely further modulate these mechanisms, potentially fostering a plastic epigenome.

### Defective *de novo* chromatin assembly in S-phase activates p53 and leads to a G0 arrest in daughter cells

CAF-1 has pleiotropic effects in S-phase cells within a few hours. Therefore, we asked how these effects impinge on cell proliferation. To do so, we used our time-lapse imaging data of the CDK2 activity sensor (Figure 2A-D) and quantified the number of cells committed to cell cycle re-entry after CAF-1 depletion. We noticed that acute CAF-1 depletion reduced the number of cells committed to enter a new cell cycle after mitosis (Figure 7A-B). This effect was stronger for cells that lost CAF-1 earlier in the cell cycle (i.e. G1 and early S-phase) compared to cells losing CAF-1 in late S or G2 phase (Figure 7B). Moreover, the vast majority of cells treated in late S or G2-phase underwent two mitoses before withdrawing from the cell cycle (i.e., G0 arrest) (Figure 7C). Thus, the time spent without CAF-1 in the cell cycle inversely correlates to the commitment of daughter cells to enter a new cell cycle. This suggests that cells need to undergo a full S-phase with defective *de novo* chromatin assembly before entering a G0 arrest. Flow cytometry measurements further validated a G0 arrest after long-term CAF-1 depletion (Supplemental Figure S7A). In addition, our single-cell transcriptomic data after 30hr CAF-1 depletion showed an increase in a G0 transcriptional state^78^ (Figure 7D, Supplemental Figure S7C and Table S1). Taken together, our results show that loss of CAF-1 precludes re-entry into a new cell cycle after experiencing a perturbed S-phase.

**Figure 7.**
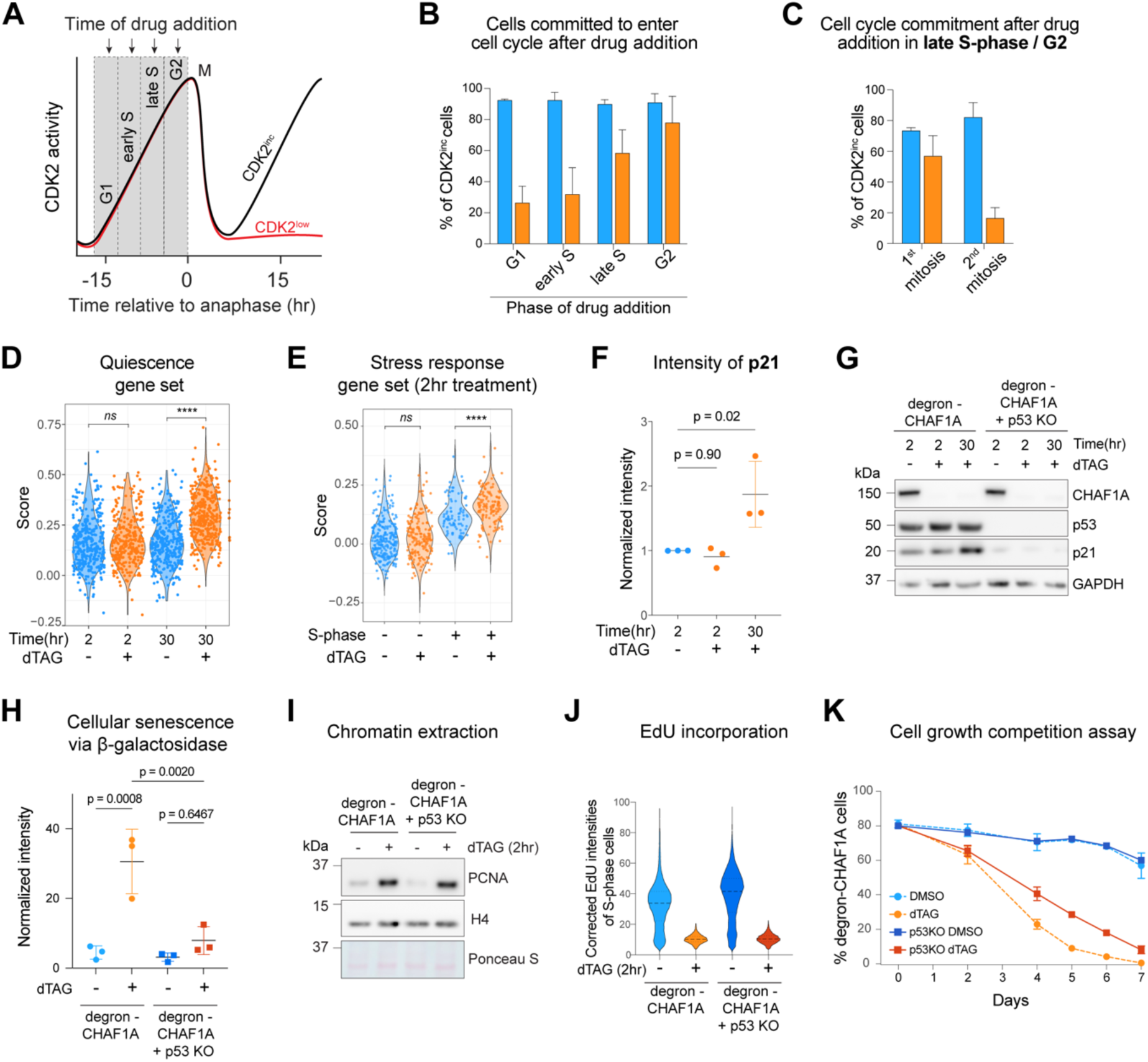
Defective *de novo* chromatin assembly in S-phase leads to a G0 arrest in daughter cells. A) Schematic representation of live-microscopy data shown in Figure 2. CDK2 increasing cells (CDK2^inc^) and CDK2 low cells (CDK2^low^) are used to calculate the percentage of cells that are committed to enter the cell cycle (CDK^inc^) vs the percentage of cells that enters G0 arrest (CDK^low^). B) Quantification of CDK2^inc^ cells of experiments shown in Supplemental Figure S2A-D. Blue = DMSO, Orange = dTAG. Error bars represent the standard deviation. C) Quantification of CDK2^inc^ cells of experiments shown in Supplemental Figure S2B and S2D separated by first and second mitosis after treatment. Blue = DMSO, Orange = dTAG. Error bars represent the standard deviation. D) Violin plot of the Quiescence score (y-axis) for DMSO or dTAG treated degron-CHAF1A cells split by 2hr and 30hrs (x-axis). Adjusted p-values were determined by Pairwise T-test and corrected Bonferroni multiple testing correction. See Table S1. E) Single cell expression analysis for stress response using ModuleScore function from Seurat for 2hr DMSO or dTAG treated CHAF1A-degron cells split by non S-phase and S-phase clusters. Adjusted p-values were determined by Pairwise T-test and corrected Bonferroni multiple testing correction. See Table S1. F) Flow cytometry analysis of p21 levels in degron-CHAF1A cells (clone 8) at indicated time points. G) Western blot of RIPA extracted cell lysates of degron-CHAF1A cells (clone 8) and degron-CHAF1A cells + p53KO at indicated time points. H) Flow cytometry analysis of cellular senescence via b-galactosidase hydrolysis. Cells were treated for 7 days with DMSO or dTAG. Intensity of CellEvent™ Senescence Green per cell was normalized to an unstained control. P-values were calculated using Students T-test. I) Western blot analysis of chromatin extracted degron-CHAF1A cells (clone 8) and degron-CHAF1A cells + p53KO after 2hr of DMSO or dTAG treatment. J) Flow cytometry analysis of EdU incorporation levels after 2hr of DMSO or dTAG treatment. Intensity of S-phase cells was corrected by mean EdU levels of corresponding G1 phase population. K) Cell growth competition assay showing proliferation efficiency in DMSO or dTAG. Degron-CHAF1A cells (clone 8) and degron-CHAF1A + p53KO cells were mixed with mNeon-tubulin hTERT-RPE-1 cells at a ratio of 80/20 at day 0. Percentage of CHAF1A-tagged cells was measured by flow cytometry as number of mNeon negative cells at indicated time points.

Interestingly, by analyzing our scRNA-seq data after 2hr of CAF-1 depletion, we find a transcriptional signature linked to a p53-dependent stress response in non-transformed cells, caused by low histone levels^79^ (Figure 7E and Table S1). Therefore, p53-dependent mechanisms could be involved in the G0 arrest. To test this hypothesis, we determined p21 proteins levels after 30hr of CAF-1 depletion. We observed that p21 was stabilized (Figure 7F), while phosphorylation of the DNA damage checkpoint kinase CHK1 remained undetected (Supplemental Figure S7B). These findings indicate that the p53-p21 axis is engaged rapidly during defective *de novo* chromatin assembly, without detectable activation of the DNA damage response. To functionally address the role of p53, we generated a TP53 knockout (p53-KO) RPE-1 degron-CHAF1A line using CRISPR genome editing. Using this line, we confirmed that p21 stabilization upon CAF-1 degradation is dependent on p53 (Figure 7G), as is entry into senescence upon longer depletions (Figure 7H). These results indicate that p53 controls the cell cycle arrest upon CAF-1 loss. Importantly, CAF-1 depleted p53-KO cells still show PCNA accumulation on chromatin (Figure 7I) and lower EdU incorporation levels (Figure 7J), which suggests that p53 does not control these CAF-1-dependent DNA replication phenotypes (Figure 1). Moreover, the absence of p53 only partially rescues proliferation defects observed following CAF-1 depletion (Figure 7K). We conclude that p53 plays a role in the early response to defective *de novo* chromatin assembly during S-phase, but additional mechanisms are likely participating in this response. It is tempting to speculate that the slowdown of DNA replication and the changes in PCNA dynamics are components of these mechanisms.

## DISCUSSION

Our work reveals the rapid multi-level response to defective *de novo* chromatin assembly during DNA replication in human cells with unprecedented temporal resolution. CAF-1 loss not only alters nascent chromatin assembly, but also immediately slows down DNA replication speed, without affecting replication timing nor triggering a canonical DNA damage checkpoint. In turn, S-phase duration is prolonged and the replicative histones supply is downregulated. In these conditions, due to perturbations in histone repertoire and defective chromatin assembly mechanisms, chromatin composition changes globally. The H3.3 variant levels increase. Chromatin maturation is delayed, with parental histones having a reduced relative abundance of H4K20me2 within a single S-phase, and transcriptional fidelity being compromised. Ultimately, these multi-level events lead to a G0 arrest that partially depends on p53. Our study uncovers mechanisms that control epigenome stability in the wake of DNA replication forks.

### Chromatin controls DNA replication speed independently of cell cycle signaling

Our work shows that CAF-1 controls DNA replication speed at a genome-wide level. This is already apparent within the shortest experimentally possible time of CAF-1 depletion, i.e. 10 minutes before EdU labeling. Thus, we propose a direct link between *de novo* chromatin assembly on nascent DNA and the speed of replication forks, as previously suggested^21^. In line with this work, PCNA plays a role in this mechanism, accumulating on chromatin together with its unloader ATAD5. Interestingly, depletion of ATAD5 phenocopies the acute loss of CAF-1, also leading to slower replication speed and an S-phase delay^56,80^. Thus, PCNA levels on chromatin control replisome progression, with CAF-1 regulating these levels. As we find plenty of PCNA in the soluble pool, we conclude that small changes in PCNA levels on chromatin can cause or be a consequence of impaired DNA replication. Since PCNA accumulates also upon perturbation of other components of the *de novo* histone deposition pathway ^21^, we favor a model where nucleosome assembly is the signal for PCNA unloading^21,80^. However, other possible mechanisms may involve the direct interaction between CAF-1 and PCNA^28,29,32^, and the binding competition to PCNA with DNA polymerases^30^. A complete understanding of these mechanisms will require further investigation.

In addition to PCNA-dependent effects, chromatin itself may hold nascent DNA strands in a conformation that favors optimal replisome progression. To support this hypothesis, we find a positive correlation between the severity of nascent chromatin opening and DNA replication slowdown in heterochromatic regions. The formal demonstration of the nature of this correlation will require development of novel technologies to understand nascent chromatin topology and organization during DNA replication^81,82^, and the understanding of the molecular mechanisms that link chromatin assembly to DNA replication. In any case, our data show that the state of newly replicated DNA has a direct influence on replisome progression genome-wide. This requires a re-thinking of the role of replicated DNA strands in controlling the progression of replication forks and their interplay with the replisome and the parental genome, also in the absence of DNA damage.

### Epigenome instability upon CAF-1 loss: different effects in constitutive and facultative heterochromatin

CAF-1 is well known for its chromatin assembly function^22,25,52^. We show that acute CAF-1 depletion affects the global composition and accessibility of chromatin within a single S-phase. While domain-scale heterochromatic PTMs distribution is not affected, we observe alterations in histone variant usage, chromatin compaction, its maturation and transcriptional fidelity within hours after CAF-1 loss. These phenotypes likely cause the epigenetic instability and plasticity that has been observed upon long-term CAF-1 depletion^20,33,36,37,55,76^. Both constitutive H3K9me3 and facultative H3K27me3 heterochromatin become hyper-accessible during DNA replication without CAF-1. Yet, these two regions differentially recover after DNA replication from this excessive opening, suggesting that local mechanisms have an impact on the consequences of defective *de novo* chromatin assembly during DNA replication.

We show that facultative heterochromatic H3K27me3 domains are particularly susceptible to CAF-1 loss, as they remain more open than the rest of the genome for several hours after DNA replication. Moreover, we find a small but significant increase in transcription of genes within these regions, suggesting a functional link between these domains and CAF-1 loss during their replication. A link had also been previously reported during *in vitro* cell differentiation^33^ and development in plants^76^. We show this likely requires replication through these domains. In *Drosophila*, Polycomb domains are sensitive to histone levels^83^, indicating a potential fragility of these regions when histones are limiting, similar to our conditions. Mechanistically, our data point to an increase of H3.3 within these domains after a single S-phase without CAF-1. In line with this, it was recently shown that the replication-dependent activity of CAF-1 acts as a mechanism to limit spurious non-canonical H3.3 deposition on chromatin in mouse oocytes and zygotes^65^. Whether H3.3 directly affects Polycomb domains by preventing local chromatin organization and modifications^84–87^, or whether H3.3 accumulation is only one aspect of a global histone imbalance, which ultimately disrupts the stability of these and other domains, remains to be investigated.

Regarding constitutive heterochromatin, it is well-established that CAF-1 binds the H3K9me3 reader protein HP1a^48,73–75^, which raised hypotheses that CAF-1 may control stability of H3K9me3 domains outside of DNA replication. In our work, H3K9me3 regions display the most severe opening during DNA replication without CAF-1. We show that these regions do not open up outside of DNA replication, implicating a role for CAF-1 in controlling accessibility of these regions during DNA replication. However, H3K9me3 domains efficiently recompact after DNA replication in a CAF-1-independent manner, and we find that they are also more compacted at steady-state levels without CAF-1. This suggests that CAF-1 loss may affect H3K9me3 heterochromatin outside the context of DNA replication. The nature of this replication-independent effect of CAF-1 remains unclear, with possible local contributions to the slowdown of replication forks in late S-phase. It will be important to shed light on these mechanisms, and the possible involvement of the *de novo* deposition of H3.3K9me3^88,89^. Indeed, we did not observe significant changes upon CAF-1 loss in H3K9me3 marks at the domain-level using ChiC-seq. We expect that higher resolution mapping of histone PTMs and their replication-coupled dynamics^90–93^ will be important approaches to further disentangle the mechanistic link between CAF-1 and heterochromatin stability.

### Towards understanding the link between CAF-1 and cell fate decisions

Our data show that the vast majority of cells that have experienced a full S-phase without CAF-1 do not continue cycling after mitosis. This underlines the essentiality of this complex^52,94^, raising important questions on the mechanisms at the core of cell fate changes upon long-term CAF-1 depletion^33,36,37,55,95,96^. Cells need to escape the cell cycle arrest to support those changes. This will require rapid adaptations. The loss of epigenome stability that we observe already within a single S-phase likely promotes the plasticity required to acquire these adaptations and to evade cell cycle arrest, with the contribution of environmental cues^20^. Understanding the nature of the adaptations that allow bypass of the cell cycle arrest is of great interest. These will reveal both the mechanisms that underpin CAF-1-mediated cell cycle control and the ones that foster cell fate change.

## Acknowledgements

We thank Kyungjae Myung for providing the ATAD5 antibodies and protocols, Bas van Steensel for sharing unpublished datasets used for initial genomics analyses, Kathleen R. Stewart-Morgan and Leonie Kollenstart for discussions on the repli-ATAC-seq setup. We thank Juan Garaycoechea, Puck Knipscheer and Jop Kind for feedback on the manuscript. We thank Anke Sparmann (Life Science Editors) for scientific editing of the manuscript. F.M. is funded by an ERC StG (851564) and the Dutch Cancer Society (2014-6649). A.v.O. is funded by European Research Council Advanced grant (ERC-AdG 101053581-scTranslatomics), Nederlandse Organisatie voor Wetenschappelijk Onderzoek (NWO) TOP award (NWO-CW 714.016.001). C.A., C.S., and S.L.S were funded by a Pew-Stewart Scholar for Cancer Research Award, an American Cancer Society Research Scholar Grant (RSG-18-008-01), and an NIH Director’s New Innovator Award (1DP2CA238330-01) to S.L.S. C.A. was also supported by NIH training grant T32 (GM065103-16). I.K.M is funded by NWO (VI.Veni.212.052). A.G. is funded by the Novo Nordisk Foundation (NNF21OC0067425), and the Lundbeck Foundation (R313-2019-448). The Novo Nordisk Foundation Center for Protein Research (CPR) is supported financially by the Novo Nordisk Foundation (grant NNF14CC0001). S.G.M. was funded by AIRC-MSCA iCARE2 fellowship, 800924, MSCA-IF 838555, and currently funded by Next generation EU-MUR MSCA Young Researcher.

## Author contributions

J.D., G.R., F.M. conceived the project. J.D., G.R., J.v.d.B, F.M. designed experiments. J.D., G.R., J.v.d.B, performed and analyzed experiments, V.B., J.v.d.B, G.R., V.v.B. M.A.V. performed bioinformatic analyses, J.F., I.K.M., R.M., J.C. performed experiments, C.A., C.S., performed live-microscopy and FISH under S.L.S. supervision, C.B., S.d.S., performed mass spectrometry under M.V.A supervision. S.G.M. shared preliminary data. I.K.M., A.B., A.G. and A.v.O. provided advice. S.L.S, A.v.O. and F.M. funded the research. F.M., J.D., G.R., J.v.d.B, wrote the manuscript with inputs from all authors, J.D., G.R., J.v.d.B. contributed equally and they are listed in alphabetic order.

## Statements (conflict of interest, diversity/inclusivity)

A.G. is co-founder and chief scientific officer (CSO) of Ankrin Therapeutics. A.G. is a member of the scientific advisory board of Molecular Cell. C.B., S.d.S. and M.A.V. are employees of MOLEQLAR Analytics GmbH.

We support inclusive, diverse, and equitable conduct of research.

## Data and code availability

The proteomics data has been deposited in the PRIDE database with accession code PXD050618. The repli-ATAC-seq, scVASA-seq and scEdU-seq data are available at GEO: GSE262518.

## STAR METHODS

### Resource availability

Lead contact: Further information and requests for resources and reagents should be directed to and will be fulfilled by the lead contact Francesca Mattiroli.

### Experimental model and subject details

Human TERT-RPE-1 cells (female) were cultured in Dulbecco’s Modified Eagle’s Medium/Nutrient Mixture F-12, GlutaMAX™ supplement (DMEM, Gibco) containing 10% Fetal Bovine Serum (FBS, Sigma Aldrich) and 100 U/mL penicillin/streptomycin (Gibco) at 37°C in 5% C02. Cells were passaged twice a week using TrypLE Express (Gibco) and tested regularly for the absence of mycoplasma.

### Method details

#### Genome editing

FKBP12^F36V^-CHAF1A expressing cells were generated by nucleofection of WT hTERT-RPE-1 cells with a CRISPR knock-in donor vector as well as a Cas9 guideRNA vector. 3×10^5 cells were resuspended in 100 uL ice cold nucleofection buffer (20 mM HEPES pH 7.5, 5 mM KCl, 10 mM MgCl2, 90 mM Na2HPO4) and 5 ug of both CRISPR-Cas9 vectors. Nucleofection was performed using Electroporation Cuvettes Plus (BTX) and the Amaxa Nucleofector II (Lonza). One nucleofection with only the Cas9 guideRNA vector served as a negative control. Cells were seeded into DMEM and left to recover for 48hr. Then, cells were selected for two weeks with 8 ug/mL blasticidin (Invivogen) before sorted as single cells into 96-well plates (BD FACSAria Fusion). For genotyping, cells of individual clones were washed with PBS0 twice before being lysed with 2-fold diluted DirectPCR Lysis Reagent (Viagen Biotech) supplemented with 20 ug/uL Proteinase K (NEB). The lysates were incubated for 16hr at 55°C followed by 1.5hr at 85°C. The resulting gDNA was genotyped using GoTaq G2 Flexi DNA Polymerase kit (Promega) and indicated primers.

For this manuscript, we used two different clonal cell lines (clone 6 and clone 8) expressing two slightly different versions of the degron-tag (Supplemental Figure S1A).

Cells stably expressing fluorescent proteins for live-microscopy experiments were generated by lentiviral transduction. For virus generation, HEK293T cells were transfected with CSII-EF DHB-mVenus and CSII-EF H2B-mTurquoise along with the packaging and envelope plasmids (pMDLg, pRSV-Rev, pCMV-VSV-G) using Fugene-HD (Promega E2311). Lentivirus was harvested 48hr after transfection, filtered through a 0.45 mm filter (Millipore), and incubated with target cells for 10hr with 5 mg/mL polybrene (EMD Millipore #TR-1003). Cells with stable integration were sorted on an Aria Fusion Flow Cytometer to establish a population expressing the fluorescent proteins.

#### Cell synchronization

To induce a G1 phase arrest, cells were cultured in DMEM medium containing 2.5% FBS and 150 nM palbociclib (Selleck Chemicals) for 24hr. For the synchronous release into S-phase, cells were washed twice with PBS0 before cultured in DMEM containing 15% FBS. To induce G2/M phase arrest, cells were treated with 300 nM nocodazole (Sigma-Aldrich) for 16 hours.

#### Drug treatments

Cells were treated with 500 nM or 1 uM dTAG^V^-1 (Bio-Techne) for indicated periods to induce degradation of FKBP12^F36V^-CHAF1A fusion protein.

#### Western Blotting

##### Whole cell extracts

Cells were harvested using TrypLE, washed once with PBS0 and pellets were lysed in RIPA buffer (25 mM Tris, 150 mM NaCl, 1% IGEPAL CA-630, 1% NaDOC, 0.1% SDS, 1x protease inhibitor and 0.5 mM DTT). The lysates were incubated on ice for 20 minutes before centrifuged at 17,000 x g for 15 minutes at 4°C. Supernatant was taken as sample for whole cell extracts. Note, when blotting for phospho-proteins, RIPA was further supplemented with 1x PhosSTOP (#4906845001 Sigma-Aldrich).

##### Separation of soluble and chromatin fractions

Cells were harvested using TrypLE, washed once with PBS0 and pellets were lysed in fractionation buffer (50 mM HEPES pH 7.9, 300 mM NaCl, 0.2 mM EDTA, 0.5% IGEPAL CA-630, 5% glycerol, 1x protease inhibitor). Lysates were centrifuged at 2800 x g for 3 minutes at 4°C and the supernatant was used as soluble fraction. The remaining chromatin pellet was washed once in fractionation buffer before being resuspended in fractionation buffer + 1 mM MgCl2 + 37.5 Units benzonase-nuclease (Merck). The chromatin pellets were incubated at 37°C for 1 hr shaking at 1000 rpm and resuspended frequently. Finally, the insoluble material was spun down at 17,000 x g for 3 minutes at 4°C and the supernatant was used as chromatin fraction.

##### Acidic extraction of histones

Cells were harvested using TrypLE, washed once with PBS0 and pellets were incubated in hypotonic buffer (10 mM HEPES pH 7.4, 10 mM KCl, 0.05% IGEPAL CA-630, 1x protease inhibitor) on ice for 10 minutes. Next, samples were spun down at 2000 x g for 5 minutes at 4°C, supernatant was removed and the pellet was washed once with hypotonic buffer. The remaining pellet was incubated in 0.2 M HCl for 45 minutes before being centrifuged at 17,000 x g for 15 minutes at 4°C. The supernatant was used as acidic extracted fraction after being neutralized in neutralization buffer (0.4 M TRIS HCL pH 8, 200 mM NaCl, 10 mM MgCl2, 1x protease inhibitor).

##### Gel electrophoresis and incubation with antibodies

Cell extracts were run on 4-12% Bolt™ Bis-Tris Plus Mini Protein Gels (ThermoScientific) and separated proteins were transferred to a 0.2 µm nitrocellulose membrane (BioRad). After blocking in 5% skimmed milk in PBS, membranes were incubated with primary antibodies in PBS and 1% skimmed milk on a roller overnight at 4°C using the concentrations indicated in the key resource table. For blotting of phospho-proteins, membranes were blocked in 2% BSA (Sigma-Aldrich) and primary antibodies were diluted in PBS and 1% BSA. After incubation with primary antibodies, membranes were washed 3x with PBST and incubated with HRP-conjugated secondary antibody (1:10,000) for 1 hour on a roller at room temperature. Next, membranes were washed 3x with PBST before signal detection using SuperSignal™ West Pico PLUS Chemiluminescent Substrate (ThermoScientific) on the Amersham ImageQuant 800 biomolecular imager.

#### Flow cytometry

Cells were labeled with EdU (10uM) for 30 minutes. Subsequently, cells were trypsinized, washed with PBS0 and fixed with 4% paraformaldehyde + 0.1% Triton-X at room temperature (RT) for 15 minutes. Next, cells were washed 2x with FACS buffer (PBS0 + 1% FBS + 0.5 mM EDTA) before being incubated in Click-IT reaction Master Mix (40mM TRIS pH 8, 4mM CuSO4*5H2O, 10mM ascorbic acid, 60 uM AF647-Azide) at RT in the dark for 45 minutes. Cells were washed with FACS buffer + 0.1% Triton-X and incubated with primary antibody at RT in the dark for 60 minutes (1/200 for anti-yH2AX, anti-pCHK1 and anti-p21). Samples were washed with FACS buffer and incubated with secondary antibody (1:500 goat-anti-rabbit AF 555) + 1ug/mL DAPI at 4°C O/N rotating. Finally, samples were washed 1x with FACS buffer + 0.1% Triton-X and 2x with FACS buffer. Samples were analyzed on BD LSR Fortessa X20 and results were analyzed using FlowJo™ v10.6.1 Software (BD Life Sciences).

#### Repli-ATAC-seq and ATAC-seq

##### Sample preparation

RPE1 cells were treated for 2 hours with 1 µM of dTAG-v1 molecule (Bio-Techne). For nascent samples, the cells were then pulsed for 20 minutes with 20 µM thymidine analog 5-ethynyl-2′-deoxyuridine (EdU, Abcam) followed by immediate harvest to assess nascent chromatin accessibility. 10×10^5 RPE1 cells per condition were counted and pooled together with 500 S2 *D.melanogaster* cells to be used for spike-in normalization. The samples were then either de-chromatinized to check for EdU incorporation (controls in Supplemental Figure S5B), or processed according to the repli-ATAC-seq protocol^97^. Post-replication samples were directly pulsed 20 minutes with 20 µM EdU after which 1 µM of dTAG-v1 molecule and 10 µM thymidine were both added for 2 hours (chase) followed by cells harvesting. Mature 2hr and 15hr samples were prepared by treating RPE1 for 2 hours with 1 µM of dTAG-v1 molecule, pulse for 20 minutes with 20 µM of EdU and then subjected to a 2- or 15-hours chase with 10 µM thymidine prior harvesting. For the de-chromatinized control samples, after cell harvesting and spike-in addition, were resuspended in 15mM Tris HCl pH 7.5 buffer complemented with 1mM EDTA and 0.5% SDS. These input samples originate from the same tube as the repli-ATAC-seq samples to ensure proper comparison. ProteinaseK (NEB) was added to the cell suspension to a final concentration of 75µg/ml followed by O.N. incubation on the shaking heat block at 55 C° 1000 rpm. The digested DNA was precipitated with 0.7 volumes of isopropanol and subjected to Tn5 (homemade) digestion after which repli-ATAC-seq samples preparation protocol was followed. To prepare classic ATAC-samples, degron-CHAF1A cells were treated with DMSO or 1 uM dTAG for 2 hrs, then 100.000 cells were mixed with 500 EdU-unlabelled *Drosophila* cells. Cell pellet was lysed in 50 µl of ATAC-resuspension buffer 1(ATAC-RSB1 10mM Tris-HCl pH 7.4, 10mM NaCl, 3mM MgCl2, Digitoxin 0.01%, Tween-20 0.1%, IGEPAL CA-630 0.1%) on ice for 3 minutes and wash out with 1 ml of ATAC-RSB2 (10mM Tris-HCl pH 7.4, 10mM NaCl, 3mM MgCl2,Tween-20 0.1%). Nuclei were subjected to Tn5 digestion (TDE1 Illumina kit) for 30 minutes at 37C° in a thermocycler shaking at 1200 rpm. DNA was cleaned up using MinElute Reaction Cleanup kit (Qiagen). Library generation was performed as for repli-ATAC-seq samples, a part from second size selection step where we used a 1.6:1 final ratio of AMPure beads (Beckman Coulter).

##### Data Analysis

Repli-ATAC libraries were sequenced 50 bp pair end modality on Illumina NextSeq2000. Raw, paired-end .fastq files from repli-ATAC and ATAC protocols were mapped to the human genome (GRCh38, Gencode) and the *Drosophila* genome (dm6, flybase), using the DNA-mapping workflow from snakePipes^98^, with the following command:

DNA-mapping --fastqc --trim --trimmer trimgalore --dedup --mapq 30 --properPairs --insertSizeMax 2000 --alignerOpts ‘ --very-sensitive’ -i <fastq_dir> -o <output_dir> -j 30

<genome>

The workflow applies paired-end trimming using trim-galore (v0.6.5) with options ‘--paired --stringency 3’, maps the reads using bowtie2 (v2.3)^99^ with options ‘--very-sensitive’, filters reads with mapq > 30 and insert size < 2000 using samtools (v1.10)^100^ and removes PCR duplicates using sambamba (v0.7.1)^101^. Further QC and creation of bigwig files was done using deepTools (v3.1)^102^.

In order to create spike-in normalized coverage files, we counted the total, de-duplicated reads mapped to the human and drosophila genomes per sample and calculated the size factor (S) per sample as follows.

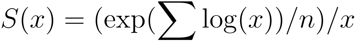

Where *x* is the counts for spike-in (drosophila) reads per sample, and refers to the number of samples in the same spike-in batch. We used this scale factor to create bigwigs using deepTools bamCoverage and bamCompare, with options ‘--binSize 15000 --normalizeUsing None --scaleFactor <scalefactor>’.

Customed R scripts were used to further analyzed Bam and Bigwig files as well as to analyze all the repli-ATAC and ATAC data. We used our ChIC-seq (DMSO control G1 sample) data as a reference to create a merged dataset with information about accessibility and genomic location about H3K9me3 and H3K27me3 heterochromatin region. This new dataset was also overlapped with the VASA-seq data to analyze changes in gene expression in the regions of interest. See Table S1.

#### scEdU-seq

scEdU-seq was performed as previously described by van den Berg et al.,^49^. Briefly, RPE-1 cells were labeled with two 15 min pulses of EdU (10uM) interspersed by 60 minutes washout with full medium. Subsequently, RPE-1 cells were trypsinized and fixed in 70% Ethanol for 24 hours at −20C. Samples were resuspended and washed in Wash Buffer (47.5mL H2O RNAse free, 1ml 1M HEPES pH 7.5, 1.5ml 5M NaCl, 3.6ul pure spermidine solution with extra 0.05% Tween, 4 ul/ml 0.5M EDTA). Subsequently, Biotin-PEG3-Azide was conjugated to EdU molecules by a CuAAC click reaction and stained with DAPI. Single S-phase RPE-1 hTERT cells were sorted into 384-well plates for scEdU-seq processing. Following the sort, libraries were prepared by the following steps; proteinaseK digestion, NlaIII genome digestion, DNA blunt ending, A-tailing and finally adapter ligation with cell barcodes and unique molecular identifiers (UMI). Single cell libraries were pooled and bound with MyOneC1 Streptavidin beads to capture DNA replication fragments. Fragments were released by heat denaturation and fragments were filled in by the Klenow enzyme. Libraries were amplified by IVT, RT and PCR and subjected to Illumina sequencing (Nextseq1000 P3 2×100bp). Code for analysis and plotting can be found on GitHub. Published Repli-Seq^103^ and EdU-seqHU^104^ data were used as reference for replication timing analysis.

#### scVASA-seq

scVASA-seq was performed as previously described by Salmen et al., 2022^58^. Briefly, total RNA is fragmented and polyadenylated. Total RNA is converted to cDNA and amplified by in vitro transcription. Ribosomal RNA sequences were depleted from Amplified RNA (aRNA) and residual RNA is converted to DNA by reverse transcription. Subsequently, libraries are amplified by PCR and subjected to Illumina sequencing (Nextseq1000 P3 2×100bp). Code for analysis and plotting can be found on GitHub.

#### ChiC-seq

ChIC-seq was performed as described previously by Zeller et al, 2023 Nature Genetics^77^. Briefly, cells were treated with DMSO or dTAG for indicated time periods. Subsequently, cells were fixed in 70% Ethanol and within 24 hours resuspended in Wash buffer (47.5ml H2O RNAse free, 1ml 1M HEPES pH 7.5, 1.5ml 5M NaCl, 3.6ul pure spermidine solution, 0.05% Tween, 200uL 0.5M EDTA & Protease inhibitor tablet). Antibodies were incubated overnight at 4°C on a roller to prevent aggegration. After washing away the antibody, pA-MNase was added to the samples in combination with DAPI. Next, a thousand cells were sorted in individual tubes per sample and Ca2+ was added to activate the pA-MNase for 30minutes at 4°C, MNase was stopped by the addition of EGTA and digested with proteinase K. The resulting DNA fragments were end-repaired, A-tailed and ligated with double-strand DNA adapters containing a T7 and UMI. Samples were amplified with a round of in vitro transcription, reverse transcriptions and finally PCR. Libraries were subjected to Illumina Sequencing (Nextseq P3 2×100bp). Code for analysis and plotting can be found on GitHub.

#### Microscopy

##### Live-cell imaging

Cells were plated on a glass bottom 96-well plate (Cellvis Cat. No. P96-1.5H-N) coated with collagen diluted 1:50 in water (Advanced BioMatrix, #5005). Cells were plated at a density of 1,200 cells per well 24hr prior to the start of imaging. For the duration of the movie, cells were maintained in at 37°C, with 5% CO2. Exposure times were set to 70ms for CFP, corresponding to H2B-mTurquoise2 and 100ms for YFP, corresponding to DHB-mVenus. After 24hr of live-cell imaging, cells were treated with either 1µM dTAG-V1 or 0.1% DMSO, followed by an additional 48hr of imaging.

##### Tracking of live-cell imaging

Live-cell tracking was conducted by generating a nuclear mask based on segmentation of the H2B-mTurquoise fluorescence signal for each frame of the live-cell movie and nearest neighbor calculations were used to track single cells. The nuclear mask was applied to the DHB-mVenus fluorescence channel and the nuclear DHB signal was measured as the mean signal of the pixels in each nucleus. Cytoplasmic DHB signal was determined by dilating the nuclear mask by 2 pixels and calculating the mean pixel intensity of the cytoplasmic ring. CDK2 activity was calculated as the ratio of cytoplasmic to nuclear DHB signal. The tracking code is available at https://github.com/scappell/Cell_tracking.

#### RNA FISH

Cells were fixed with 4% paraformaldehyde for 15min. Histone H4.2 (Thermo Fisher, VA6-3174283-VC) mRNA was visualized according to the manufacturer’s protocol (ViewRNA ISH Cell Assay Kit, ThermoFisher QVC0001). Cells were permeabilized for 30min and mRNA probes hybridized for 4hr at 40°C. Exposure time was set to 600ms for Cy5. Histone H4.2 mRNA FISH signal was quantified as the median pixel value of a two-pixel-wide cytoplasmic ring around the nuclear mask.

#### Quantitative Mass spectrometry

##### Sample preparation for histone modification analysis by MS

Approximately 500k RPE-1 degron-CHAF1A cells were synchronized in G2 phase as described in Figure 3A and under method section “cell synchronization”. Acid extracted histones were processed according to a SP3 protocol as described previously^105^. However, proprietary steps developed by MOLEQLAR Analytics GmbH have been added in order to adjust the protocol for histone specific aspects. Upon overnight digestion at 37°C and 1,000rpm in a table-top thermomixer, samples were acidified by adding 5µl of 5% TFA and quickly vortexed. Beads were immobilized on a magnetic rack, and peptides were recovered by transferring the supernatant to new tubes. Samples were dried down using a vacuum concentrator and reconstituted by adding 12µl 0.1% FA to reach a peptide concentration of approximately 0.2µg/µl. MS injection-ready samples were stored at −20°C.

##### LC-MS analysis of histone modifications

Approximately 200ng of peptides from each sample were separated on a C18 column (Aurora Elite TS, 15 cm x 75 μm ID, 1.7 μm, IonOpticks) with a gradient from 5% B to 25% B (solvent A 0.1% FA in water, solvent B 100% ACN, 0.1% FA) over 35min at a flow rate of 300nl/min (Vanquish Neo UHPLC-Systems, Thermo - Fisher, San Jose, CA) and directly sprayed into a Exploris 240 mass spectrometer (Thermo-Fisher Scientific). The mass spectrometer was operated in full-scan mode to identify and quantify specific fragment ions of N-terminal peptides of human histone 3.1 and histone 4 proteins. Survey full scan MS spectra (from m/z 250–1600) were acquired with resolution 60,000 at m/z 400 (AGC target of 3×106). Typical mass spectrometric conditions were: spray voltage, 1.9kV; no sheath and auxiliary gas flow; heated capillary temperature, 300°C.

##### Quantification of histone modifications

Data analysis was performed with Skyline (version 22.2.0.351)^106^ by using doubly and triply charged peptide masses for extracted ion chromatograms (XICs). Peaks were selected manually. Heavy arginine-labelled spiketides (13C6; 15N4) were used to confirm the correct retention times. The SILAC peptides (i.e. heavy arginine-labelled peptides (R6 Arginine 13C6)) were selected to assess the newly heavy arginine incorporation into synthetized histones proteins. To that aim, we removed any peptides that did not contain any arginine. Integrated peak values (Total Area MS1) under the first 4 isotopomers were used for further calculations and log2 transformed. Since the three G1 samples are unlabeled, we used them to evaluate the labeling quality, defined as our ability to not detect hits to peptides containing heavy arginine in G1 condition and measured as the median of 100-100 x intensitySILAC/(intensitySILAC+intensitylight) percentage over the peptides equal to 98.7, 99.4, and 99.5% for replicates 1, 2, 3 respectively. Subsequently, any SILAC signal in G1 samples was not considered (i.e. set to 0).

For a given precursor, the percentage of each modification on the following H3/H4 lysines: K4, K9, K14, K18, K23 K27, K36, K56, K79/K5, K8, K12, K16 (all denoted as H4_4…17), K20 within the same peptide is derived from the ratio of this structural modified peptide intensity to the sum of all isotopically similar peptides intensities using either SILAC (new) or light intensities (old). In other words, the Total Area MS1 value was used to calculate the relative abundance of an observed modified peptide as percentage of the overall peptide. The intensity of the unmodified peptide of histone 3.1 (aa 41–49) was used as a proxy of total histone 3.1. Coeluting isobaric modifications were quantified using three unique MS2 fragment ions.

Per modification, we tested for any significant difference in differential relative abundance between old and new histones in DMSO versus dTAG condition (i.e. effect of dTAG) using a t-test under the following null hypothesis (rel. abund. from light – rel. abund. from SILAC) DMSO = (rel. abund. from light – rel. abund. from SILAC) dTAG. Benjamini-Hochberg multiple-testing procedure has then been applied.

#### Analysis of senescence-associated β-galactosidase activity

For quantification of SA-b-gal activity by flow cytometry, the CellEvent™ Senescence Green Flow Cytometry Assay Kit (ThermoScientific) was used. Cells were treated with either DMSO or dTAG (500 nM) for 7 days without splitting. Next, cells were harvested by trypsinization, washed once with PBS0 and then fixed with 4% paraformaldehyde for 10min at RT. Subsequently, samples were washed with PBS0 and incubated for 2hr in 1:750 CellEventTM Senescence Green Probe diluted in CellEventTM Senescence Buffer at 37°C shaking (1000 rpm). Lastly, cells were washed with PBS0 and analyzed using BD LSR Fortessa X20 (BD Biosciences).

#### Cell growth competition assay

RPE-1 mNeonGreen-tagged *β*-tubulin cells were seeded at a 1:5 ratio with either degron-CHAF1A cells or p53KO degron-CHAF1A cells. Cells were treated with dTAG (500 nM) or DMSO over a period of 7 days and split at day 0, 2 and 4 according to their confluency. The remaining cells of every timepoint were collected for flow cytometry analysis, washed 1x in FACS buffer and the percentage of degron-CHAF1A cells was assessed as mNeon negative population using BD LSR Fortessa X20 (BD Biosciences).

## SUPPLEMENTAL FILES

### SUPPLEMENTAL FIGURES

**Supplemental Figure S1.**
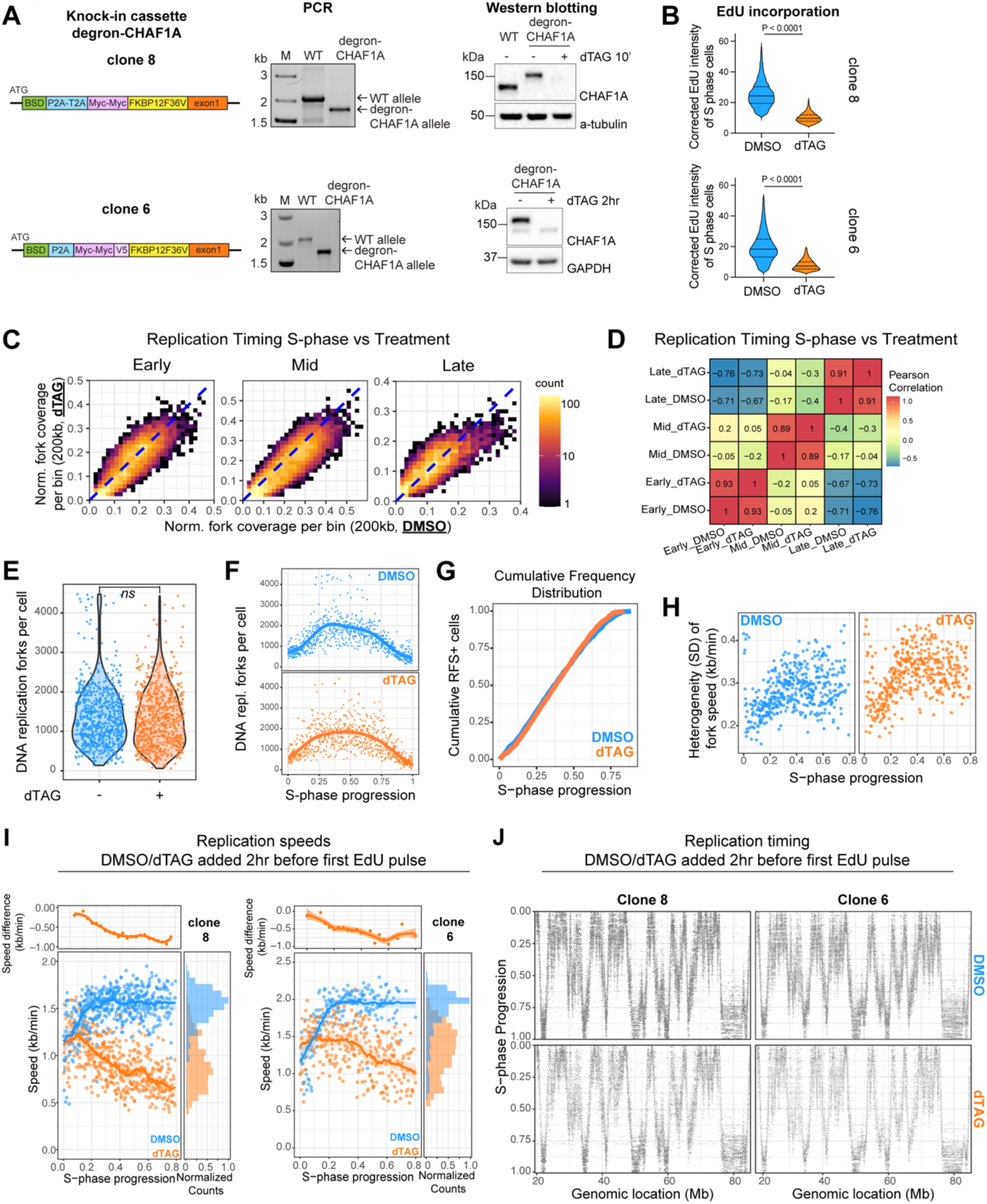
Related to Figure 1. A) Left: Overview of the knock-in constructs used to tag the N-terminus of the CHAF1A subunit. Middle: PCR gel showing insertion of the knock-in constructs into the genome of hTERT-RPE1 cells. Right: Western blot analysis of degron-CHAF1A clones treated with DMSO or dTAG for indicated time periods. B) Violin plots of EdU incorporation levels of S-phase cells measured by flow cytometry. EdU intensity in S-phase cells was normalized to the intensity in G1 cells. C) Bivariate heatmap of the average number of replication forks per bin per cell for DMSO (x-axis) compared to dTAG (y-axis) treated degron-CHAF1A cells split by early, middle and late S-phase pseudo-bulk samples (color indicates counts per bin). D) Pearson correlation matrix representation for each pseudo-bulk and degron-CHAF1A treatment (DMSO or dTAG), number and colors indicate Pearson correlation. E) Violin plots of the number of DNA replication forks per single cell in degron-CHAF1A cells treated with DMSO or dTAG. Adjusted p-values were obtained by Pairwise T-test followed by Bonferroni multiple testing correction. F) The number of DNA replication forks (y-axis) per single cell in degron-CHAF1A cells treated with DMSO (blue) or dTAG (orange) over S-phase progression (x-axis). G) The Cumulative frequency distribution (y-axis) of cells with a replication fork speed (RFS+) cells over scEdU-seq+ cells over S-phase Progression (x-axis). H) Heterogeneity (SD) of DNA replication speeds within single cells (y-axis) over S-phase progression (x-axis). I) DNA replication speed over S-phase in degron-CHAF1A (clone 8 and clone 6) treated with DMSO (blue) or dTAG (orange) for 2hr followed by scEdUseq. Difference in DNA replication speeds between DMSO and dTAG in kb/min (y-axis) over S-phase progression (x-axis, top-left), marginal density (x-axis) of DNA replication speed in kb/min (y-axis) colored for DMSO- (blue) or dTAG-treated cells (orange, bottom-right). J) Heatmap showing scEdU-seq maximum normalized log counts for 2-hours DMSO-treated and dTAG-treated RPE-1 degron-CHAF1A cells (as in panel I) ordered according to S-phase progression (y-axis) and binned per 40 kb bins (x-axis) for a 50 megabase region of chromosome 2.

**Supplemental Figure S2.**
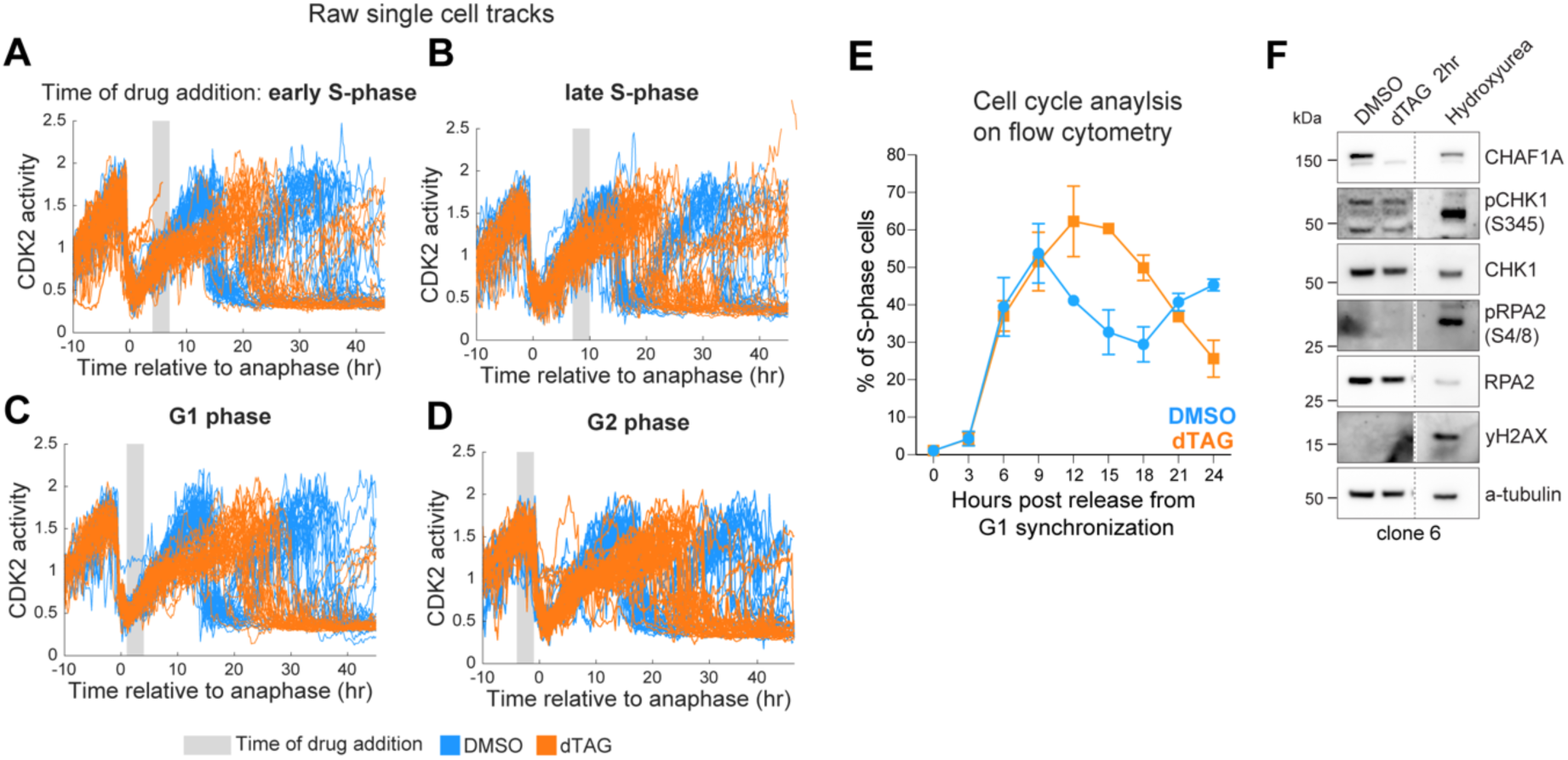
Related to Figure 2. A) -D) Single-cell traces of CDK2 activity aligned computationally to the time of anaphase. Cells included in the different analyses experienced the start of DMSO (blue) or dTAG (orange, 1uM) treatment relative to their last anaphase in indicated cell cycle phases. E) Flow cytometry analysis of percentage of S-phase cells based on EdU/DAPI staining (y-axis). Cells were arrested in G1 phase by treatment with CDK4/6 inhibitor palbociclib (150nM for 24hr). One hour before release into S-phase, DMSO or dTAG was added to the medium. Cells were harvested every 3hr post release (x-axis). Error bars represent SD. F) Western blot analysis of phosphorylation of Chk1, RPA and H2AX phosphorylation in degron-CHAF1A clone 6 cells treated with DMSO, 2hr dTAG and hydroxyurea (2mM, 24hr), probed with indicated antibodies.

**Supplemental Figure S3.**
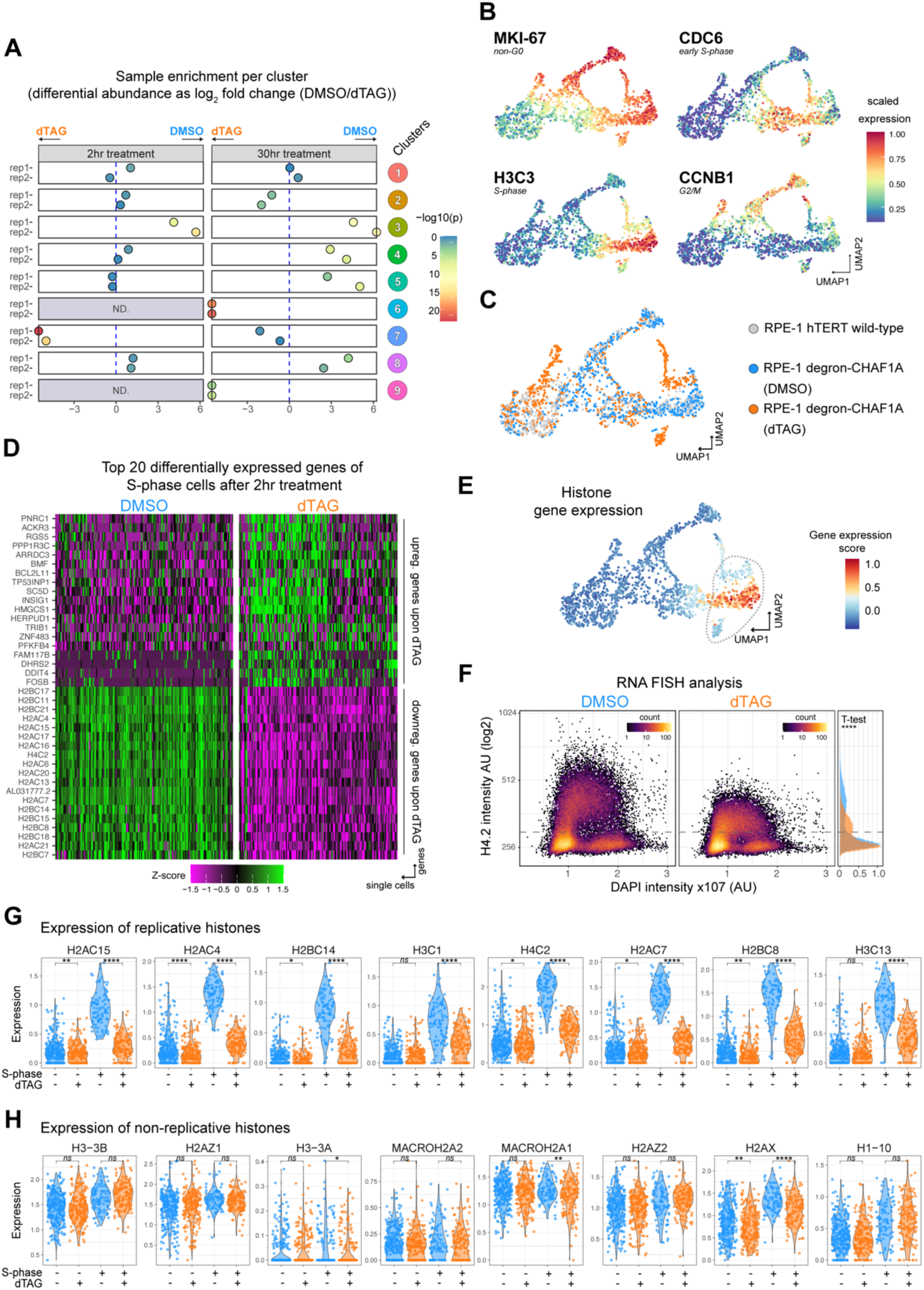
Related to Figure 3. A) Differential abundance (log_2_) of cells treated with DMSO or dTAG for 2 or 30hr per Leiden Cluster, colors indicate −log10(adj. p-value) for the Binomial Test. The value of the datapoints that are drawn on the borders of the graph is infinite. B) UMAPs determined using total RNA counts per gene illustrating scaled expression for MKI-67, CDC6, H3C3 and CCNB1 in all scVASA-seq cells. C) UMAPs determined using total RNA counts per gene illustrating different cell lines and treatments. D) Heatmap of expression (Z-scored by gene) for the top 20 differentially expressed genes (y-axis) split by DMSO and 2hr dTAG treatment for degron-CHAF1A cells (x-axis) in S-phase cells E) UMAPs determined using total RNA counts per gene illustrating Histone Gene expression (see Table S1) after 2hr treatment using the ModuleScore function in Seurat. F) Quantification of smFISH for H4 (y-axis) versus DAPI intensity (x-axis) in degron-CHAF1A degron cells treated with DMSO and dTAG (2hr). G) Violin plots of individual replicative histone gene expression (y-axis) in indicated conditions (x-axis), where each dot represents an individual cell. Adjusted p-values were determined by Pairwise T-test and corrected Bonferroni multiple testing correction. H) Violin plots of individual non-replicative histone gene expression (y-axis) in indicated conditions (x-axis), where each dot represents an individual cell. Adjusted p-values were determined by Pairwise T-test and corrected Bonferroni multiple testing correction.

**Supplemental Figure S4.**
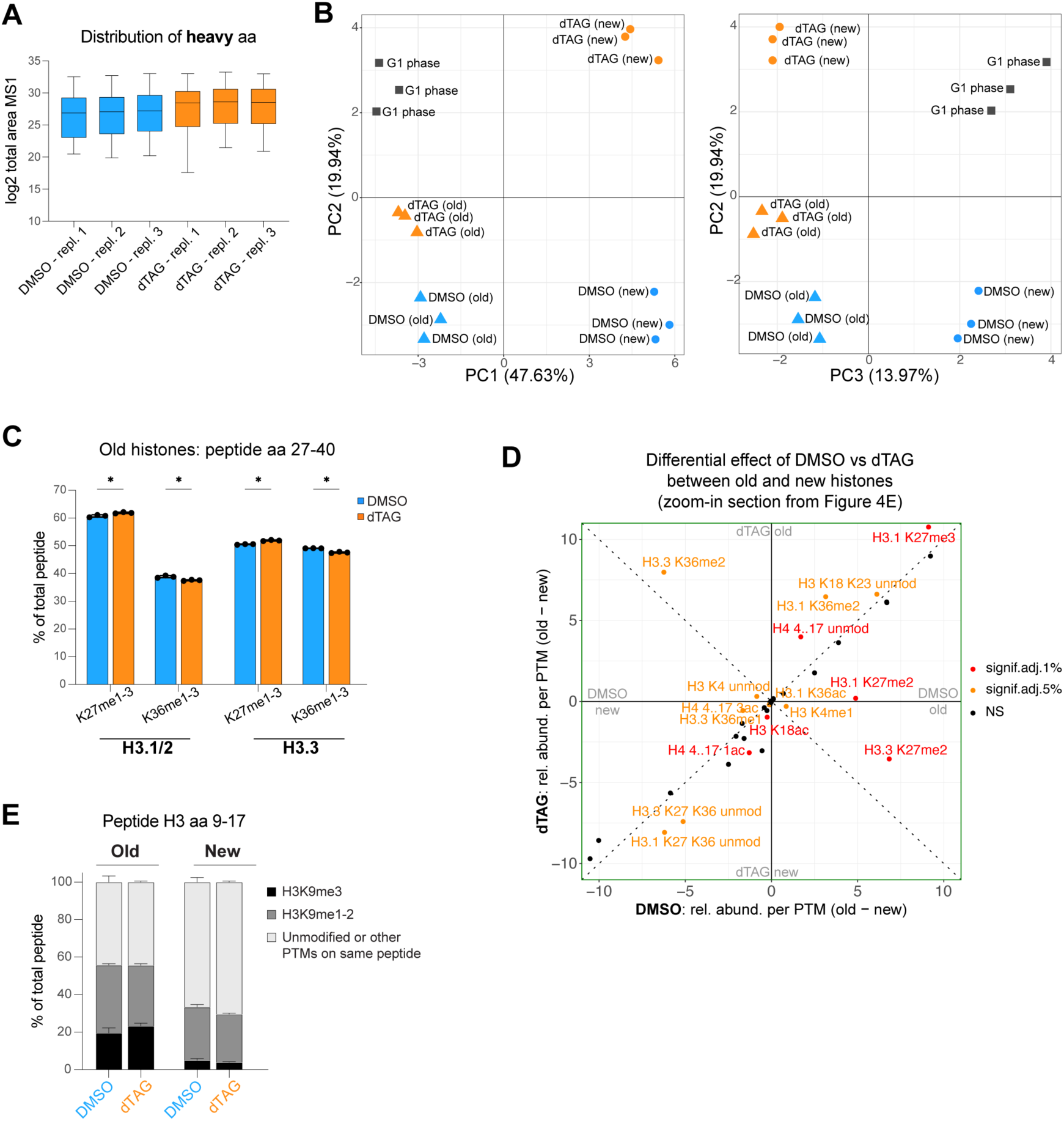
Related to Figure 4. A) Boxplots showing distribution of peptide-wise heavy log2 intensities per sample. B) Principal component analysis from the PTMs relative abundance of new and old histones across conditions allows to visualize clustering of samples. C) Quantification of old H3.1/2 and H3.3 peptides (amino acids 27-40). Relative abundances of mono-, di- and tri-methylation of K27 or K36 were summed up. D) Zoom-in of Figure 4E. E) Quantification of H3K9-anchored PTMs relative abundance on old and new H3 peptide 9-17 associated with DMSO or dTAG treatment.

**Supplemental Figure S5.**
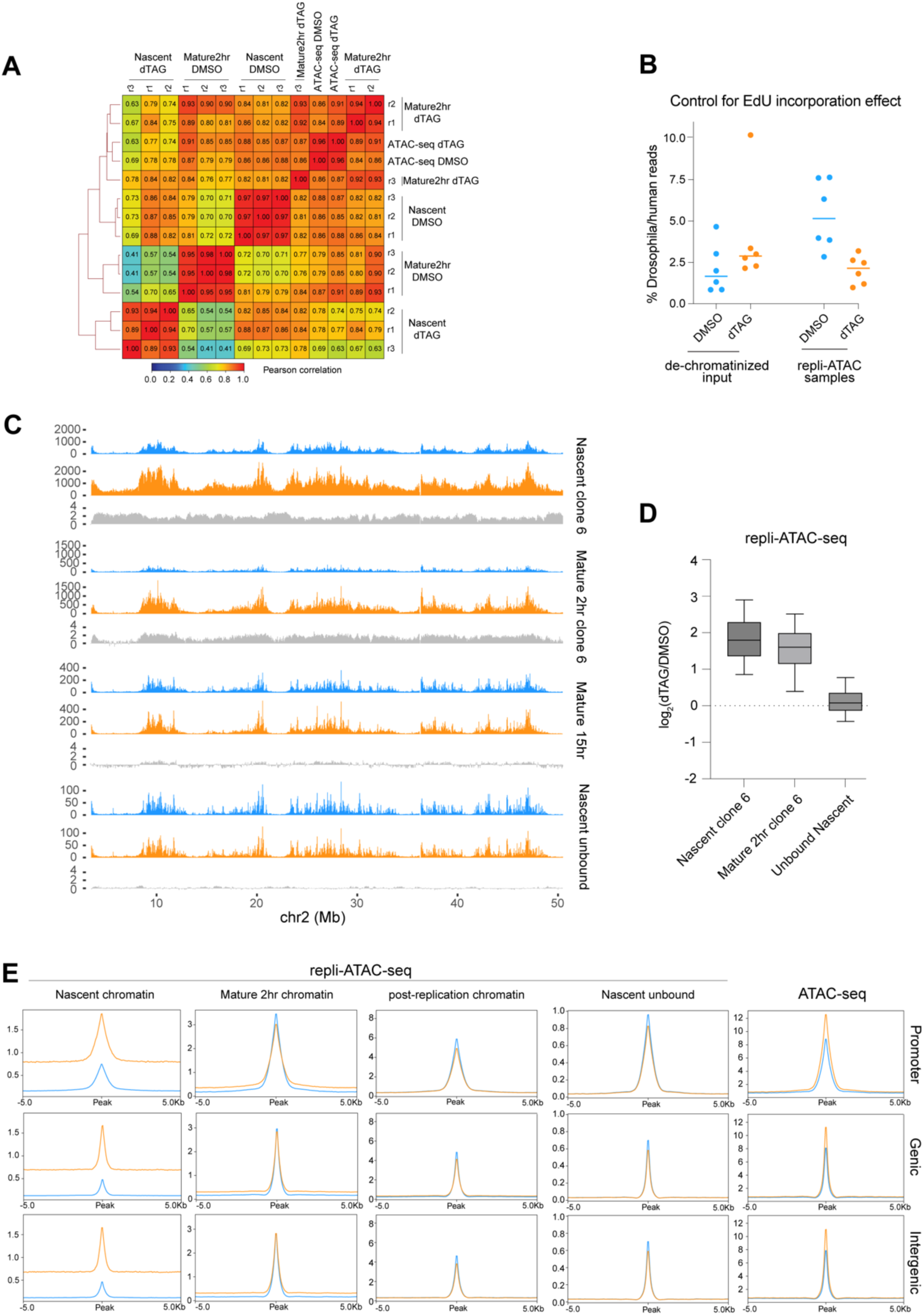
Related to Figure 5. A) Pearson correlation matrix of repli-ATAC-seq and ATAC-seq samples described in this manuscript. The dendrogram on the left of the heatmap represents the hierarchical clustering of the rows. Numbers and colors indicate Pearson correlation value B) Ratio between spiked-in *Drosophila melanogaster* and human EdU reads of proteinaseK pretreated (de-chromatinized DNA samples) and repli-ATAC-seq samples. Horizontal bars represent median values. These data are used to monitor the effects of EdU incorporation in processed repli-ATAC-seq samples. C) Spike-in normalized signal (y-axis) of nascent and mature 2hr chromatin repli-ATAC-seq samples from clone 6 (top) and 15hr mature chromatin repli-ATAC-seq sample from clone 8 (middle). Bottom panel shows RPM-normalized unbound (EdU unlabeled DNA) nascent chromatin repli-ATAC-seq sample from clone 8. The signal is visualized over a selected region on chromosome 2. In blue DMSO control samples, in orange dTAG treated samples, in grey Log_2_FC quantification of dTAG over DMSO signal. D) Boxplot showing accessibility changes of dTAG over DMSO signal for nascent and mature 2hr repli-ATAC-seq samples in clone 6 and unbound repli-ATAC-seq samples in clone 8. E) Comparison of average accessibility signal (y-axis) from nascent, mature, post-replication chromatin and nascent unbound repli-ATAC-seq and ATAC-seq samples within ±5Kb regions around annotated peaks in genes promoters, gene bodies and intergenic regions.

**Supplemental Figure S6.**
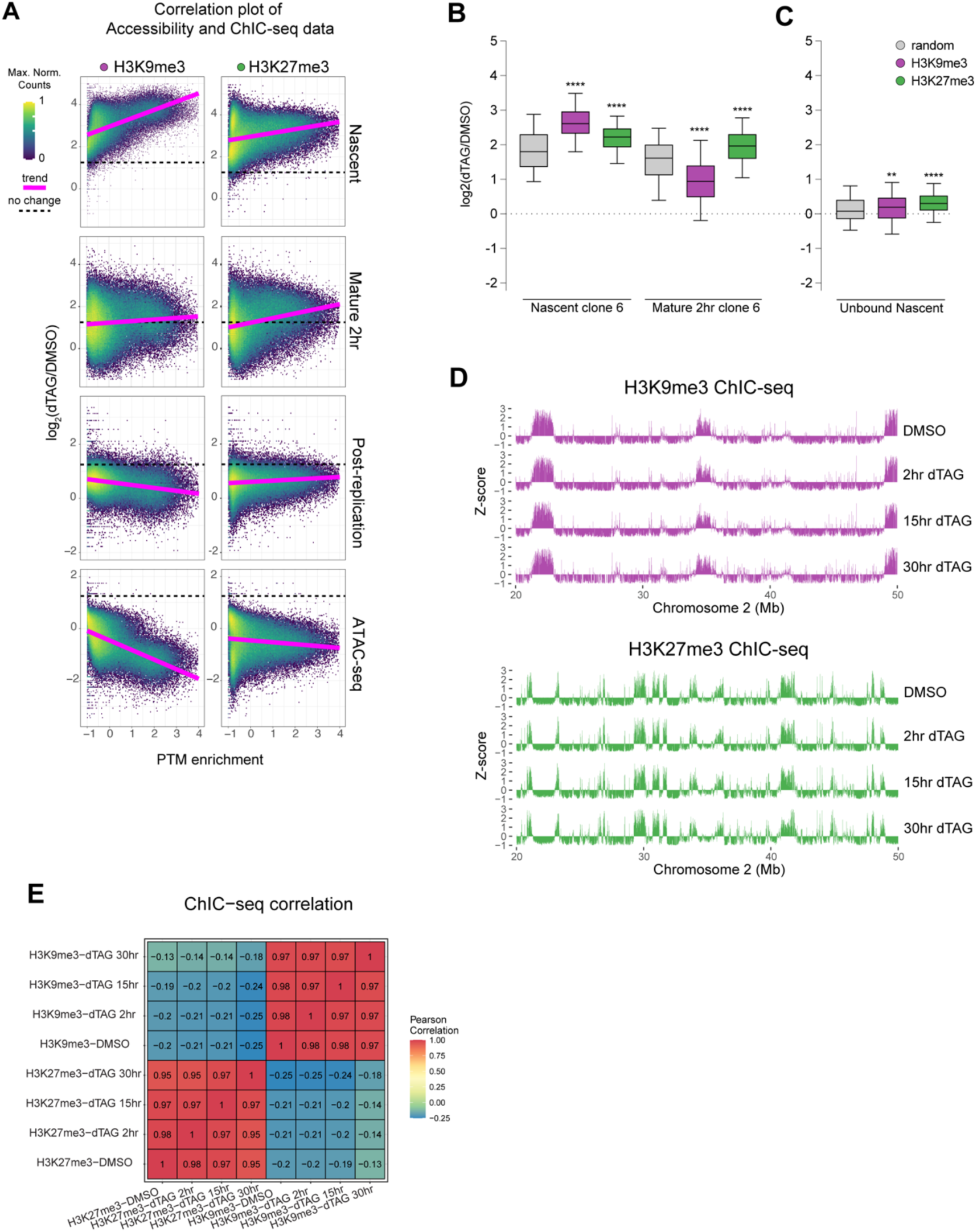
Related to Figure 6. A) Scatterplot showing correlation of chromatin accessibility Log_2_(dTAG/DMSO) signal (y-axis) over ChIC-seq signal from DMSO samples (x-axis) per 15kb bin along the genome. Calculated trend lines are shown in magenta. B-C) Boxplot quantifying change in chromatin accessibility signal upon CAF-1 removal, in heterochromatin regions in the spike-in-normalized Nascent and Mature (2hr) sample for clone 6 (B) and unbound samples from nascent clone 8 (C). The top 1000 bins have been used for calculation as in Figure 6 B-C. One-way ANOVA test was performed to determine statistically significant differences between each genomic region. D) Z-score ChIC-seq signal representative of H3K9me3 and H3K27me3 marks in untreated DMSO and dTAG at different time points (rows) over a selected region of 30 Mb on chromosome 2. E) Pearson correlation matrix of all ChIC-seq samples.

**Supplemental Figure S7.**
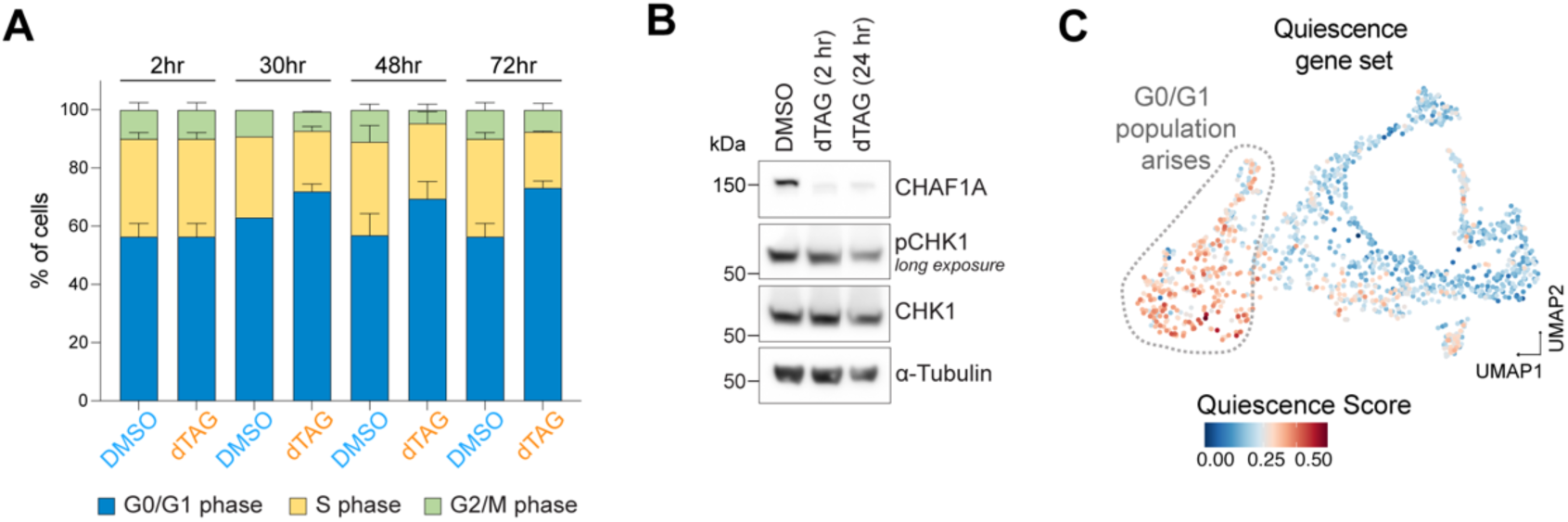
Related to Figure 7. A) Cumulative barplots of cell cycle distribution of unsynchronized degron-CHAF1A cells (clone 8) after treatment with DMSO or dTAG for indicated time periods. Cells were pulsed with EdU for 30min before harvest and cell cycle distribution was analyzed by flow cytometry using DAPI/EdU intensities. B) Western blot analysis of RIPA extracted degron-CHAF1A cells (clone 8) after DMSO and dTAG treatment for indicated time points. C) Same UMAP plot as Figure 3B with cells colored by Quiescence score (see Table S1) obtained with Seurat’s ModuleScore function.

### SUPPLEMENTAL TABLES and FILES

**Supplemental Table S1. Lists of gene sets**

List of gene sets that were used to calculate expression differences in transcription based on scVASA seq data set

**Supplemental Table S2. SILAC-MS on histone variants and their PTMs**

Intensity of light peptides aggregated by modification by precursor per sample
Intensity of heavy peptides aggregated by modification by precursor per sample
PTM relative abundance in old and new histones (grouped per precursor peptide)

**Supplemental File**

Script repli-ATAC-seq.txt

## Notes

### Summary of Updates

We updated funding information, and phrasing throughout the manuscript.

## REFERENCES

1. Yadav, T., Quivy, J.-P., and Almouzni, G. (2018). Chromatin plasticity: A versatile landscape that underlies cell fate and identity. Science (1979) 361, 1332–1336. 10.1126/science.aat8950.

2. Millán-Zambrano, G., Burton, A., Bannister, A.J., and Schneider, R. (2022). Histone post-translational modifications — cause and consequence of genome function. Nat Rev Genet 23, 563–580. 10.1038/s41576-022-00468-7.

3. Escobar, T.M., Loyola, A., and Reinberg, D. (2021). Parental nucleosome segregation and the inheritance of cellular identity. Nat Rev Genet. 10.1038/s41576-020-00312-w.

4. Stewart-Morgan, K.R., Petryk, N., and Groth, A. (2020). Chromatin replication and epigenetic cell memory. Nat Cell Biol 22. 10.1038/s41556-020-0487-y.

5. Du, W., Shi, G., Shan, C.-M., Li, Z., Zhu, B., Jia, S., Li, Q., and Zhang, Z. (2022). Mechanisms of chromatin-based epigenetic inheritance. Sci China Life Sci 65, 2162– 2190. 10.1007/s11427-022-2120-1.

6. Ye, X., and Adams, P.D. (2003). Coordination of S-Phase Events and Genome Stability. Cell Cycle 2, 184–186. 10.4161/cc.2.3.389.

7. MacAlpine, D.M., and Almouzni, G. (2013). Chromatin and DNA Replication. Cold Spring Harb Perspect Biol 5, a010207–a010207. 10.1101/cshperspect.a010207.

8. Mendiratta, S., Gatto, A., and Almouzni, G. (2019). Histone supply: Multitiered regulation ensures chromatin dynamics throughout the cell cycle. Journal of Cell Biology 218. 10.1083/jcb.201807179.

9. Alabert, C., Jasencakova, Z., and Groth, A. (2017). Chromatin replication and histone dynamics. Adv Exp Med Biol 1042, 311–333. 10.1007/978-981-10-6955-0_15.

10. Hammond, C.M., Strømme, C.B., Huang, H., Patel, D.J., and Groth, A. (2017). Histone chaperone networks shaping chromatin function. Nat Rev Mol Cell Biol 18, 141–158. 10.1038/nrm.2016.159.

11. Alabert, C., and Groth, A. (2012). Chromatin replication and epigenome maintenance. Nat Rev Mol Cell Biol 13, 153–167. 10.1038/nrm3288.

12. Alabert, C., Barth, T.K., Reverón-Gómez, N., Sidoli, S., Schmidt, A., Jensen, O.N., Imhof, A., and Groth, A. (2015). Two distinct modes for propagation of histone PTMs across the cell cycle. Genes Dev 29, 585–590. 10.1101/gad.256354.114.

13. Bandau, S., Alvarez, V., Jiang, H., Graff, S., Sundaramoorthy, R., Gierlinski, M., Toman, M., Owen-Hughes, T., Sidoli, S., Lamond, A., et al. (2024). RNA polymerase II promotes the organization of chromatin following DNA replication. EMBO Rep. 10.1038/s44319-024-00085-x.

14. Alvarez, V., Bandau, S., Jiang, H., Rios-Szwed, D., Hukelmann, J., Garcia-Wilson, E., Wiechens, N., Griesser, E., Ten Have, S., Owen-Hughes, T., et al. (2023). Proteomic profiling reveals distinct phases to the restoration of chromatin following DNA replication. Cell Rep 42, 111996. 10.1016/j.celrep.2023.111996.

15. Groth, A., Corpet, A., Cook, A.J.L., Roche, D., Bartek, J., Lukas, J., and Almouzni, G. (2007). Regulation of replication fork progression through histone supply and demand. Science (1979) 318, 1928–1931. 10.1126/science.1148992.

16. Duronio, R.J., and Marzluff, W.F. (2017). Coordinating cell cycle-regulated histone gene expression through assembly and function of the Histone Locus Body. RNA Biol 14, 726–738. 10.1080/15476286.2016.1265198.

17. Saldivar, J.C., Hamperl, S., Bocek, M.J., Chung, M., Bass, T.E., Cisneros-Soberanis, F., Samejima, K., Xie, L., Paulson, J.R., Earnshaw, W.C., et al. (2018). An intrinsic S/G _2_ checkpoint enforced by ATR. Science (1979) 361, 806–810. 10.1126/science.aap9346.

18. Ye, X., Franco, A.A., Santos, H., Nelson, D.M., Kaufman, P.D., and Adams, P.D. (2003). Defective S phase chromatin assembly causes DNA damage, activation of the S phase checkpoint, and S phase arrest. Mol Cell 11, 341–351. 10.1016/S1097-2765(03)00037-6.

19. Marzluff, W.F., and Duronio, R.J. (2002). Histone mRNA expression: Multiple levels of cell cycle regulation and important developmental consequences. Curr Opin Cell Biol 14, 692–699. 10.1016/S0955-0674(02)00387-3.

20. Franklin, R., Murn, J., and Cheloufi, S. (2021). Cell Fate Decisions in the Wake of Histone H3 Deposition. Front Cell Dev Biol 9. 10.3389/fcell.2021.654915.

21. Mejlvang, J., Feng, Y., Alabert, C., Neelsen, K.J., Jasencakova, Z., Zhao, X., Lees, M., Sandelin, A., Pasero, P., Lopes, M., et al. (2014). New histone supply regulates replication fork speed and PCNA unloading. Journal of Cell Biology 204, 29–43. 10.1083/jcb.201305017.

22. Smith, S., and Stillman, B. (1989). Purification and characterization of CAF-I, a human cell factor required for chromatin assembly during DNA replication in vitro. Cell 58, 15–25. 10.1016/0092-8674(89)90398-X.

23. Smith, S., and Stillman, B. (1991). Stepwise assembly of chromatin during DNA replication in vitro. EMBO J 10, 971–980.

24. Kaufman, P.D., Kobayashi, R., Kessler, N., and Stillman, B. (1995). The p150 and p60 subunits of chromatin assemblyfactor I: A molecular link between newly synthesized histories and DNA replication. Cell 81, 1105–1114. 10.1016/S0092-8674(05)80015-7.

25. Verreault, A., Kaufman, P.D., Kobayashi, R., and Stillman, B. (1996). Nucleosome assembly by a complex of CAF-1 and acetylated histones H3/H4. Cell 87, 95–104. 10.1016/S0092-8674(00)81326-4.

26. Kamakaka, R.T., Bulger, M., Kaufman, P.D., Stillman, B., and Kadonaga, J.T. (1996). Postreplicative chromatin assembly by Drosophila and human chromatin assembly factor 1. Mol Cell Biol 16, 810–817. 10.1128/MCB.16.3.810.

27. Tyler, J.K., Adams, C.R., Chen, S.-R., Kobayashi, R., Kamakaka, R.T., and Kadonaga, J.T. (1999). The RCAF complex mediates chromatin assembly during DNA replication and repair. Nature 402, 555–560. 10.1038/990147.

28. Shibahara, K.I., and Stillman, B. (1999). Replication-dependent marking of DNA by PCNA facilitates CAF-1-coupled inheritance of chromatin. Cell 96, 575–585. 10.1016/S0092-8674(00)80661-3.

29. Moggs, J.G., Grandi, P., Quivy, J.-P., Jonsson, Z.O., Hubscher, U., Becker, P.B., and Almouzni, G. (2000). A CAF-1-PCNA-Mediated Chromatin Assembly Pathway Triggered by Sensing DNA Damage. Mol Cell Biol 20, 1206–1218. 10.1128/MCB.20.4.1206-1218.2000.

30. Rouillon, C., Eckhardt, B. V, Kollenstart, L., Gruss, F., Verkennis, A.E.E., Rondeel, I., Krijger, P.H.L., Ricci, G., Biran, A., van Laar, T., et al. (2023). CAF-1 deposits newly synthesized histones during DNA replication using distinct mechanisms on the leading and lagging strands. Nucleic Acids Res. 10.1093/nar/gkad171.

31. Ouasti, F., Audin, M., Fréon, K., Quivy, J.-P., Tachekort, M., Cesard, E., Thureau, A., Ropars, V., Fernández Varela, P., Moal, G., et al. (2024). Disordered regions and folded modules in CAF-1 promote histone deposition in Schizosaccharomyces pombe. Elife 12. 10.7554/eLife.91461.

32. Zhang, Z., Shibahara, K.I., and Stillman, B. (2000). PCNA connects DNA replication to epigenetic inheritance in yeast. Nature 408, 221–225. 10.1038/35041601.

33. Cheng, L., Zhang, X., Wang, Y., Gan, H., Xu, X., Lv, X., Hua, X., Que, J., Ordog, T., and Zhang, Z. (2019). Chromatin Assembly Factor 1 (CAF-1) facilitates the establishment of facultative heterochromatin during pluripotency exit. Nucleic Acids Res. 10.1093/nar/gkz858.

34. Krawitz, D.C., Kama, T., and Kaufman, P.D. (2002). Chromatin assembly factor I mutants defective for PCNA binding require Asf1/Hir proteins for silencing. Mol Cell Biol 22, 614–625. 10.1128/MCB.22.2.614.

35. Ben-Shahar, T.R., Castillo, A.G., Osborne, M.J., Borden, K.L.B., Kornblatt, J., and Verreault, A. (2009). Two Fundamentally Distinct PCNA Interaction Peptides Contribute to Chromatin Assembly Factor 1 Function. Mol Cell Biol 29, 6353–6365. 10.1128/MCB.01051-09.

36. Cheloufi, S., Elling, U., Hopfgartner, B., Jung, Y.L., Murn, J., Ninova, M., Hubmann, M., Badeaux, A.I., Euong Ang, C., Tenen, D., et al. (2015). The histone chaperone CAF-1 safeguards somatic cell identity. Nature 528, 218–224. 10.1038/nature15749.

37. Ishiuchi, T., Enriquez-Gasca, R., Mizutani, E., Boškoviä, A., Ziegler-Birling, C., Rodriguez-Terrones, D., Wakayama, T., Vaquerizas, J.M., and Torres-Padilla, M.E. (2015). Early embryonic-like cells are induced by downregulating replication-dependent chromatin assembly. Nat Struct Mol Biol 22, 662–671. 10.1038/nsmb.3066.

38. Ridgway, P., and Almouzni, G. (2000). CAF-1 and the inheritance of chromatin states: at the crossroads of DNA replication and repair. J Cell Sci 113 (Pt 1), 2647–2658.

39. Almouzni, G., and Wolffe, A.P. (1993). Replication-coupled chromatin assembly is required for the repression of basal transcription in vivo. Genes Dev 7, 2033–2047. 10.1101/gad.7.10.2033.

40. Ramachandran, S., and Henikoff, S. (2016). Transcriptional Regulators Compete with Nucleosomes Post-replication. Cell 165, 580–592. 10.1016/j.cell.2016.02.062.

41. Nakano, S., Stillman, B., and Horvitz, H.R. (2011). Replication-coupled chromatin assembly generates a neuronal bilateral asymmetry in C. elegans. Cell 147, 1525– 1536. 10.1016/j.cell.2011.11.053.

42. Nabet, B., Roberts, J.M., Buckley, D.L., Paulk, J., Dastjerdi, S., Yang, A., Leggett, A.L., Erb, M.A., Lawlor, M.A., Souza, A., et al. (2018). The dTAG system for immediate and target-specific protein degradation. Nat Chem Biol 14, 431–441. 10.1038/s41589-018-0021-8.

43. Natsume, T., Kiyomitsu, T., Saga, Y., and Kanemaki, M.T. (2016). Rapid Protein Depletion in Human Cells by Auxin-Inducible Degron Tagging with Short Homology Donors. Cell Rep 15, 210–218. 10.1016/j.celrep.2016.03.001.

44. Liu, C.-P., Yu, Z., Xiong, J., Hu, J., Song, A., Ding, D., Yu, C., Yang, N., Wang, M., Yu, J., et al. (2023). Structural insights into histone binding and nucleosome assembly by chromatin assembly factor-1. Science (1979) 381. 10.1126/science.add8673.

45. Mattiroli, F., Gu, Y., Yadav, T., Balsbaugh, J.L.J.L., Harris, M.R.M.R., Findlay, E.S.E.S., Liu, Y., Radebaugh, C.A.C.A., Stargell, L.A.L.A., Ahn, N.G.N.G., et al. (2017). DNA-mediated association of two histone-bound complexes of yeast chromatin assembly factor-1 (CAF-1) drives tetrasome assembly in the wake of DNA replication. Elife 6, e22799. 10.7554/eLife.22799.

46. Sauer, P.V., Timm, J., Liu, D., Sitbon, D., Boeri-Erba, E., Velours, C., Mücke, N., Langowski, J., Ochsenbein, F., Almouzni, G., et al. (2017). Insights into the molecular architecture and histone H3-H4 deposition mechanism of yeast chromatin assembly factor 1. Elife 6, 835–839. 10.7554/eLife.23474.

47. Liu, W.H., Roemer, S.C., Zhou, Y., Shen, Z.J., Dennehey, B.K., Balsbaugh, J.L., Liddle, J.C., Nemkov, T., Ahn, N.G., Hansen, K.C., et al. (2016). The Cac1 subunit of histone chaperone CAF-1 organizes CAF-1-H3/H4 architecture and tetramerizes histones. Elife 5, 2852–2861. 10.7554/eLife.18023.001.

48. Quivy, J.-P.P., Gérard, A., Cook, A.J.L.L., Roche, D., and Almouzni, G. (2008). The HP1-p150/CAF-1 interaction is required for pericentric heterochromatin replication and S-phase progression in mouse cells. Nat Struct Mol Biol 15, 972–979. 10.1038/nsmb.1470.

49. Berg, J. van den, Batenburg, V. van, and Oudenaarden, A. van (2022). Acceleration of genome replication uncovered by single-cell nascent DNA sequencing. bioRxiv, 2022.12.13.520365. 10.1101/2022.12.13.520365.

50. Rhind, N., and Gilbert, D.M. (2013). DNA Replication Timing. Cold Spring Harb Perspect Biol 5, a010132–a010132. 10.1101/cshperspect.a010132.

51. Trotter, E.W., and Hagan, I.M. (2020). Release from cell cycle arrest with Cdk4/6 inhibitors generates highly synchronized cell cycle progression in human cell culture. Open Biol 10. 10.1098/rsob.200200.

52. Hoek, M., and Stillman, B. (2003). Chromatin assembly factor 1 is essential and couples chromatin assembly to DNA replication in vivo. Proceedings of the National Academy of Sciences 100, 12183–12188. 10.1073/pnas.1635158100.

53. Günesdogan, U., Jäckle, H., and Herzig, A. (2014). Histone supply regulates S phase timing and cell cycle progression. Elife 3, e02443. 10.7554/eLife.02443.

54. Nabatiyan, A., and Krude, T. (2004). Silencing of chromatin assembly factor 1 in human cells leads to cell death and loss of chromatin assembly during DNA synthesis. Mol Cell Biol 24, 2853–2862. 10.1128/MCB.24.7.2853.

55. Franklin, R., Guo, Y., He, S., Chen, M., Ji, F., Zhou, X., Frankhouser, D., Do, B.T., Chiem, C., Jang, M., et al. (2022). Regulation of chromatin accessibility by the histone chaperone CAF-1 sustains lineage fidelity. Nat Commun 13, 2350. 10.1038/s41467-022-29730-6.

56. Lee, K., Fu, H., Aladjem, M.I., and Myung, K. (2013). ATAD5 regulates the lifespan of DNA replication factories by modulating PCNA level on the chromatin. Journal of Cell Biology 200, 31–44. 10.1083/jcb.201206084.

57. Bellelli, R., Belan, O., Pye, V.E., Clement, C., Maslen, S.L., Skehel, J.M., Cherepanov, P., Almouzni, G., and Boulton, S.J. (2018). POLE3-POLE4 Is a Histone H3-H4 Chaperone that Maintains Chromatin Integrity during DNA Replication. Mol Cell. 10.1016/j.molcel.2018.08.043.

58. Salmen, F., De Jonghe, J., Kaminski, T.S., Alemany, A., Parada, G.E., Verity-Legg, J., Yanagida, A., Kohler, T.N., Battich, N., van den Brekel, F., et al. (2022). High-throughput total RNA sequencing in single cells using VASA-seq. Nat Biotechnol 40, 1780–1793. 10.1038/s41587-022-01361-8.

59. Mendiratta, S., Ray-Gallet, D., Lemaire, S., Gatto, A., Forest, A., Kerlin, M.A., and Almouzni, G. (2024). Regulation of replicative histone RNA metabolism by the histone chaperone ASF1. Mol Cell. 10.1016/j.molcel.2023.12.038.

60. Nelson, D.M., Ye, X., Hall, C., Santos, H., Ma, T., Kao, G.D., Yen, T.J., Harper, J.W., and Adams, P.D. (2002). Coupling of DNA Synthesis and Histone Synthesis in S Phase Independent of Cyclin/cdk2 Activity. Mol Cell Biol 22, 7459–7472. 10.1128/MCB.22.21.7459-7472.2002.

61. Armstrong, C., Passanisi, V.J., Ashraf, H.M., and Spencer, S.L. (2023). Cyclin E/CDK2 and feedback from soluble histone protein regulate the S phase burst of histone biosynthesis. Cell Rep 42, 112768. 10.1016/j.celrep.2023.112768.

62. Völker-Albert, M.C., Schmidt, A., Barth, T.K., Forne, I., and Imhof, A. (2018). Detection of Histone Modification Dynamics during the Cell Cycle by MS-Based Proteomics. In, pp. 61–74. 10.1007/978-1-4939-8663-7_4.

63. Loyola, A., Bonaldi, T., Roche, D., Imhof, A., and Almouzni, G. (2006). PTMs on H3 Variants before Chromatin Assembly Potentiate Their Final Epigenetic State. Mol Cell 24, 309–316. 10.1016/j.molcel.2006.08.019.

64. Ray-Gallet, D., Woolfe, A., Vassias, I., Pellentz, C., Lacoste, N., Puri, A., Schultz, D.C., Pchelintsev, N.A., Adams, P.D., Jansen, L.E.T., et al. (2011). Dynamics of Histone H3 Deposition In Vivo Reveal a Nucleosome Gap-Filling Mechanism for H3.3 to Maintain Chromatin Integrity. Mol Cell 44, 928–941. 10.1016/j.molcel.2011.12.006.

65. Ishiuchi, T., Abe, S., Inoue, K., Yeung, W.K.A., Miki, Y., Ogura, A., and Sasaki, H. (2021). Reprogramming of the histone H3.3 landscape in the early mouse embryo. Nat Struct Mol Biol 28, 38–49. 10.1038/s41594-020-00521-1.

66. Alabert, C., Loos, C., Voelker-Albert, M., Graziano, S., Forné, I., Reveron-Gomez, N., Schuh, L., Hasenauer, J., Marr, C., Imhof, A., et al. (2020). Domain Model Explains Propagation Dynamics and Stability of Histone H3K27 and H3K36 Methylation Landscapes. Cell Rep 30, 1223–1234.e8. 10.1016/j.celrep.2019.12.060.

67. Saredi, G., Huang, H., Hammond, C.M., Alabert, C., Bekker-Jensen, S., Forne, I., Reverón-Gómez, N., Foster, B.M., Mlejnkova, L., Bartke, T., et al. (2016). H4K20me0 marks post-replicative chromatin and recruits the TONSL-MMS22L DNA repair complex. Nature 534, 714–718. 10.1038/nature18312.

68. Pellegrino, S., Michelena, J., Teloni, F., Imhof, R., and Altmeyer, M. (2017). Replication-Coupled Dilution of H4K20me2 Guides 53BP1 to Pre-replicative Chromatin. Cell Rep 19, 1819–1831. 10.1016/j.celrep.2017.05.016.

69. Simonetta, M., de Krijger, I., Serrat, J., Moatti, N., Fortunato, D., Hoekman, L., Bleijerveld, O.B., Altelaar, A.F.M., and Jacobs, J.J.L. (2018). H4K20me2 distinguishes pre-replicative from post-replicative chromatin to appropriately direct DNA repair pathway choice by 53BP1-RIF1-MAD2L2. Cell Cycle 17, 124–136. 10.1080/15384101.2017.1404210.

70. Scharf, A.N.D., Meier, K., Seitz, V., Kremmer, E., Brehm, A., and Imhof, A. (2009). Monomethylation of Lysine 20 on Histone H4 Facilitates Chromatin Maturation. Mol Cell Biol 29, 57–67. 10.1128/MCB.00989-08.

71. Stewart-Morgan, K.R., Reverón-Gómez, N., and Groth, A. (2019). Transcription Restart Establishes Chromatin Accessibility after DNA Replication. Mol Cell, 284–297. 10.1016/j.molcel.2019.04.033.

72. Chen, B., MacAlpine, H.K., Hartemink, A.J., and MacAlpine, D.M. (2023). Spatiotemporal kinetics of CAF-1-dependent chromatin maturation ensures transcription fidelity during S-phase. Genome Res 33, 2108–2118. 10.1101/gr.278273.123.

73. Quivy, J.P., Roche, D., Kirschner, D., Tagami, H., Nakatani, Y., and Almouzni, G. (2004). A CAF-1 dependent pool of HP1 during heterochromatin duplication. EMBO Journal 23, 3516–3526. 10.1038/sj.emboj.7600362.

74. Loyola, A., Tagami, H., Bonaldi, T., Roche, D., Quivy, J.P., Imhof, A., Nakatani, Y., Dent, S.Y.R., and Almouzni, G. (2009). The HP1α-CAF1-SetDB1-containing complex provides H3K9me1 for Suv39-mediated K9me3 in pericentric heterochromatin. EMBO Rep 10, 769–775. 10.1038/embor.2009.90.

75. Murzina, N., Verreault, A., Laue, E., and Stillman, B. (1999). Heterochromatin dynamics in mouse cells: Interaction between chromatin assembly factor 1 and HP1 proteins. Mol Cell 4, 529–540. 10.1016/S1097-2765(00)80204-X.

76. Jiang, D., and Berger, F. (2017). DNA replication–coupled histone modification maintains Polycomb gene silencing in plants. Science (1979) 357, 1146–1149. 10.1126/science.aan4965.

77. Zeller, P., Yeung, J., Viñas Gaza, H., de Barbanson, B.A., Bhardwaj, V., Florescu, M., van der Linden, R., and van Oudenaarden, A. (2023). Single-cell sortChIC identifies hierarchical chromatin dynamics during hematopoiesis. Nat Genet 55, 333–345. 10.1038/s41588-022-01260-3.

78. Riba, A., Oravecz, A., Durik, M., Jiménez, S., Alunni, V., Cerciat, M., Jung, M., Keime, C., Keyes, W.M., and Molina, N. (2022). Cell cycle gene regulation dynamics revealed by RNA velocity and deep-learning. Nat Commun 13, 2865. 10.1038/s41467-022-30545-8.

79. Sokolova, M., Turunen, M., Mortusewicz, O., Kivioja, T., Herr, P., Vähärautio, A., Björklund, M., Taipale, M., Helleday, T., and Taipale, J. (2017). Genome-wide screen of cell-cycle regulators in normal and tumor cells identifies a differential response to nucleosome depletion. Cell Cycle 16, 189–199. 10.1080/15384101.2016.1261765.

80. Kang, M.-S., Kim, J., Ryu, E., Ha, N.Y., Hwang, S., Kim, B.-G., Ra, J.S., Kim, Y.J., Hwang, J.M., Myung, K., et al. (2019). PCNA Unloading Is Negatively Regulated by BET Proteins. Cell Rep 29, 4632–4645.e5. 10.1016/j.celrep.2019.11.114.

81. Mitter, M., Gasser, C., Takacs, Z., Langer, C.C.H., Tang, W., Jessberger, G., Beales, C.T., Neuner, E., Ameres, S.L., Peters, J.-M., et al. (2020). Conformation of sister chromatids in the replicated human genome. Nature 586, 139–144. 10.1038/s41586-020-2744-4.

82. Oomen, M.E., Hedger, A.K., Watts, J.K., and Dekker, J. (2020). Detecting chromatin interactions between and along sister chromatids with SisterC. Nat Methods 17, 1002–1009. 10.1038/s41592-020-0930-9.

83. McPherson, J.-M.E., Grossmann, L.C., Salzler, H.R., Armstrong, R.L., Kwon, E., Matera, A.G., McKay, D.J., and Duronio, R.J. (2023). Reduced histone gene copy number disrupts Drosophila Polycomb function. Genetics 224. 10.1093/genetics/iyad106.

84. Blackledge, N.P., and Klose, R.J. (2021). The molecular principles of gene regulation by Polycomb repressive complexes. Nat Rev Mol Cell Biol 22, 815–833. 10.1038/s41580-021-00398-y.

85. Yu, J.-R., Lee, C.-H., Oksuz, O., Stafford, J.M., and Reinberg, D. (2019). PRC2 is high maintenance. Genes Dev 33, 903–935. 10.1101/gad.325050.119.

86. Chen, P., Zhao, J., Wang, Y., Wang, M., Long, H., Liang, D., Huang, L., Wen, Z., Li, W., Li, X., et al. (2013). H3.3 actively marks enhancers and primes gene transcription via opening higher-ordered chromatin. Genes Dev 27, 2109–2124. 10.1101/gad.222174.113.

87. Liu, C., Yu, J., Song, A., Wang, M., Hu, J., Chen, P., Zhao, J., and Li, G. (2023). Histone H1 facilitates restoration of H3K27me3 during DNA replication by chromatin compaction. Nat Commun 14, 4081. 10.1038/s41467-023-39846-y.

88. Carraro, M., Hendriks, I.A., Hammond, C.M., Solis-Mezarino, V., Völker-Albert, M., Elsborg, J.D., Weisser, M.B., Spanos, C., Montoya, G., Rappsilber, J., et al. (2023). DAXX adds a de novo H3.3K9me3 deposition pathway to the histone chaperone network. Mol Cell 83, 1075–1092.e9. 10.1016/j.molcel.2023.02.009.

89. Navarro, C., Lyu, J., Katsori, A.-M., Caridha, R., and Elsässer, S.J. (2020). An embryonic stem cell-specific heterochromatin state promotes core histone exchange in the absence of DNA accessibility. Nat Commun 11, 5095. 10.1038/s41467-020-18863-1.

90. Ostrowski, M.S., Yang, M.G., McNally, C.P., Abdulhay, N.J., Wang, S., Nora, E.P., Goodarzi, H., and Ramani, V. (2023). The single-molecule accessibility landscape of newly replicated mammalian chromatin. bioRxiv, 2023.10.09.561582. 10.1101/2023.10.09.561582.

91. Reverón-Gómez, N., González-Aguilera, C., Stewart-Morgan, K.R., Petryk, N., Flury, V., Graziano, S., Johansen, J.V., Jakobsen, J.S., Alabert, C., and Groth, A. (2018). Accurate Recycling of Parental Histones Reproduces the Histone Modification Landscape during DNA Replication. Mol Cell 72. 10.1016/j.molcel.2018.08.010.

92. Liu, Y., Zhangding, Z., Liu, X., Gan, T., Ai, C., Wu, J., Liang, H., Chen, M., Guo, Y., Lu, R., et al. (2024). Fork coupling directs DNA replication elongation and termination. Science (1979) 383, 1215–1222. 10.1126/science.adj7606.

93. Gaggioli, V., Lo, C.S.Y., Reverón-Gómez, N., Jasencakova, Z., Domenech, H., Nguyen, H., Sidoli, S., Tvardovskiy, A., Uruci, S., Slotman, J.A., et al. (2023). Dynamic de novo heterochromatin assembly and disassembly at replication forks ensures fork stability. Nat Cell Biol 25, 1017–1032. 10.1038/s41556-023-01167-z.

94. Houlard, M., Berlivet, S., Probst, A. V., Quivy, J.P., Héry, P., Almouzni, G., and Gérard, M. (2006). CAF-1 is essential for heterochromatin organization in pluripotent embryonic cells. PLoS Genet 2, 1686–1696. 10.1371/journal.pgen.0020181.

95. Replogle, J.M., Saunders, R.A., Pogson, A.N., Hussmann, J.A., Lenail, A., Guna, A., Mascibroda, L., Wagner, E.J., Adelman, K., Lithwick-Yanai, G., et al. (2022). Mapping information-rich genotype-phenotype landscapes with genome-scale Perturb-seq. Cell 185, 2559–2575.e28. 10.1016/j.cell.2022.05.013.

96. Ng, C., Aichinger, M., Nguyen, T., Au, C., Najar, T., Wu, L., Mesa, K.R., Liao, W., Quivy, J.-P., Hubert, B., et al. (2019). The histone chaperone CAF-1 cooperates with the DNA methyltransferases to maintain *Cd4* silencing in cytotoxic T cells. Genes Dev 33, 669–683. 10.1101/gad.322024.118.

97. Stewart-Morgan, K.R., and Groth, A. (2023). Profiling Chromatin Accessibility on Replicated DNA with repli-ATAC-Seq. In, pp. 71–84. 10.1007/978-1-0716-2899-7_6.

98. Bhardwaj, V., Heyne, S., Sikora, K., Rabbani, L., Rauer, M., Kilpert, F., Richter, A.S., Ryan, D.P., and Manke, T. (2019). snakePipes: facilitating flexible, scalable and integrative epigenomic analysis. Bioinformatics 35, 4757–4759. 10.1093/bioinformatics/btz436.

99. Langmead, B., and Salzberg, S.L. (2012). Fast gapped-read alignment with Bowtie 2. Nat Methods 9, 357–359. 10.1038/nmeth.1923.

100. Li, H., Handsaker, B., Wysoker, A., Fennell, T., Ruan, J., Homer, N., Marth, G., Abecasis, G., and Durbin, R. (2009). The Sequence Alignment/Map format and SAMtools. Bioinformatics 25, 2078–2079. 10.1093/bioinformatics/btp352.

101. Tarasov, A., Vilella, A.J., Cuppen, E., Nijman, I.J., and Prins, P. (2015). Sambamba: fast processing of NGS alignment formats. Bioinformatics 31, 2032–2034. 10.1093/bioinformatics/btv098.

102. Ramírez, F., Ryan, D.P., Grüning, B., Bhardwaj, V., Kilpert, F., Richter, A.S., Heyne, S., Dündar, F., and Manke, T. (2016). deepTools2: a next generation web server for deep-sequencing data analysis. Nucleic Acids Res 44, W160–W165. 10.1093/nar/gkw257.

103. Dekker, J., Belmont, A.S., Guttman, M., Leshyk, V.O., Lis, J.T., Lomvardas, S., Mirny, L.A., O’Shea, C.C., Park, P.J., Ren, B., et al. (2017). The 4D nucleome project. Nature 549, 219–226. 10.1038/nature23884.

104. Macheret, M., and Halazonetis, T.D. (2018). Intragenic origins due to short G1 phases underlie oncogene-induced DNA replication stress. Nature 555, 112–116. 10.1038/nature25507.

105. Hughes, C.S., Moggridge, S., Müller, T., Sorensen, P.H., Morin, G.B., and Krijgsveld, J. (2019). Single-pot, solid-phase-enhanced sample preparation for proteomics experiments. Nat Protoc 14, 68–85. 10.1038/s41596-018-0082-x.

106. MacLean, B., Tomazela, D.M., Shulman, N., Chambers, M., Finney, G.L., Frewen, B., Kern, R., Tabb, D.L., Liebler, D.C., and MacCoss, M.J. (2010). Skyline: an open source document editor for creating and analyzing targeted proteomics experiments. Bioinformatics 26, 966–968. 10.1093/bioinformatics/btq054.

